# Interspecific introgression and natural selection in the evolution of Japanese apricot (*Prunus mume*)

**DOI:** 10.1101/2020.06.23.141200

**Authors:** Koji Numaguchi, Takashi Akagi, Yuto Kitamura, Ryo Ishikawa, Takashige Ishii

## Abstract

- Domestication and population differentiation in crops involve considerable phenotypic changes. The logs of these evolutionary paths, including natural/artificial selection, can be found in the genomes of the current populations. However, these profiles have been little studied in tree crops, which have specific characters, such as long generation time and clonal propagation, maintaining high levels of heterozygosity.
- We conducted exon-targeted resequencing of 129 genomes in the genus *Prunus*, mainly Japanese apricot (*Prunus mume*), and apricot (*P. armeniaca*), plum (*P. salicina*), and peach (*P. persica*). Based on their genome-wide single nucleotide polymorphisms merged with published resequencing data of 79 Chinese *P. mume* cultivars, we inferred complete and ongoing population differentiation in *P. mume*.
- Sliding window characterization of the indexes for genetic differentiation identified interspecific fragment introgressions between *P. mume* and related species (plum and apricot). These regions often exhibited strong selective sweeps formed in the paths of establishment or formation of substructures of *P. mume*, suggesting that *P. mume* has frequently imported advantageous genes from other species in the subgenus *Prunus* as adaptive evolution.
- These findings shed light on the complicated nature of adaptive evolution in a tree crop that has undergone interspecific exchange of genome fragments with natural/artificial selection.

## Introduction

Domestication and population differentiation often involve considerable phenotypic changes in crops (Diamond, 2002; Purugganan & Fuller, 2009; Zeder, 2015). Mainly natural mutations are thought to drive these changes, while occasional interspecific introgression can also potentially contribute (Baack & Rieseberg, 2007; Harrison & Larson, 2014; Gaut *et al.*, 2015; Suarez-Gonzalez *et al.*, 2018). For instance, domesticated apple (*Malus domestica*), which originated from a wild apple species distributed in Central Asia (*M. sieversiii*), has recently experienced additional genomic introgression from another wild species (*M. sylvestris*) (Cornille *et al.*, 2012). Citrus and olive are also suggested to have complicated evolutionary pathways involving interspecific introgression from related or ancestral wild species (Wu *et al.*, 2014, 2018; Diez *et al.*, 2015). Hexaploid bread wheat (*Triticum aestivum*) is a notable example of herbaceous crops with a drastic domestication process. This species was developed from a dynamic hybridization between tetraploid emmer wheat (*T. turgidum*) and diploid Tausch’s goatgrass (*Aegilops tauschii*) and subsequent introgression from other species promoted cultivar differentiation (Molnár-Láng *et al.*, 2015; He *et al.*, 2019). Owing to the evolutionary importance of interspecific introgression, much previous research on a variety of species has inferred the presence of interspecific introgression in current populations. Notwithstanding, few studies have further estimated the genome-wide distribution of introgressed fragments and their importance for domestication/population differentiation events.

Introduced mutations favorable to environmental adaptation or human preference might be subjected to natural or artificial selection. When a particular locus experiences a strong selection pressure, the genetic diversity of adjacent genomic regions is reduced as well as the targeted locus itself, which is known as a “selective sweep” (Stephan, 2019). Therefore, we can estimate the genetic factors playing important roles in the formation of current populations by characterizing genome-wide selective sweep profiles (Clark *et al.*, 2004; Sabeti *et al.*, 2007; Kosova *et al.*, 2010; Ishii *et al.*, 2013; Akagi *et al.*, 2016; Lee *et al.*, 2016; Pankin *et al.*, 2018; Nadachowska-Brzyska *et al.*, 2019). Selective sweep profiles have been well studied especially in annual crops such as rice (*Oryza sativa*). In annual crops, where selected alleles are thought to be fixed in a homozygous state, patterns of selective sweeps have been identified mainly using site frequency spectrum (SFS)-based methods using the reduction in genetic diversity as the index. Conversely, perennial crops (or tree crops) have more complicated genomic/genetic conditions, mainly due to vegetative propagation, frequent outcrossing, and long generation time. Therefore, a selected allele in perennial crops is expected to be maintained in a heterozygous manner, which would be quite similar to animal (including human) genomes, requiring haplotype-based detection of selective sweeps (Voight *et al.*, 2006; Sabeti *et al.*, 2007).

The genus *Prunus* includes a wide variety of major tree crops consumed worldwide, such as peach (*P. persica*), sweet cherry (*P. avium*), plum (*P. salicina*), apricot (*P. armeniaca*), and almond (*P. dulcis*). Japanese apricot (*P. mume*) is also a major fruit/flower crop in East Asia commonly known as Chinese “mei” or “mume”. Japanese apricot is believed to have been domesticated firstly in China several thousand years ago, transferred into Japan ca. 2,000 years ago, originally for ornamental purposes (Mega *et al.*, 1988; Horiuchi *et al.*, 1996; Faust *et al.*, 2011). Currently, cultivars are widely diversified mainly based on their usage, such as for pickles (“umeboshi”), syrups/ liquors, and ornamental flowers. However, in contrast to historical implication and conventional categorization, the genetic background of this species remains little known. Japanese apricot is morphologically similar to apricot and plum, and they are all nested in the subgenus *Prunus* (Bortiri *et al.*, 2001). Species of the subgen. *Prunus* are partially compatible for interspecific crossing (Yamaguchi *et al.*, 2018; Morimoto *et al.*, 2019), and some Japanese apricot cultivars have been traditionally thought to carry genetic factors of apricot (*P. armeniaca*) (Mega *et al.*, 1988; Horiuchi *et al.*, 1996), which was later supported by molecular marker analyses (Shimada *et al.*, 1994; Hayashi *et al.*, 2008; Numaguchi *et al.*, 2019). Furthermore, recent breeding programs often utilize interspecific crossing of the subgen. *Prunus*, such as “Sumomo-ume” (*P. salicina* × *P. mume*), or “Pluot” (*P. salicina* × *P. armeniaca*) (Kyotani *et al.*, 1988; Brantley, 2004; Yaegaki *et al.*, 2012).

Given the above information, it would appear that Japanese apricot, and related species in the subgen. *Prunus*, have undergone complicated evolutionary processes, involving natural or artificial selection and potential introgressions among them. Here, to clarify the evolutionary paths to establish the current subgen. *Prunus*, mainly for *P. mume*, we analyzed genome-wide single nucleotide polymorphisms (SNPs) based on targeted resequencing of ca. 15,000 exons in East Asian *P. mume* cultivars. An integrative analysis of selective sweeps and the transition of fragmental genetic structures successfully inferred the importance of interspecific introgressions and lineage-specific selection during the evolution of *P. mume*.

## Materials and Methods

### Plant materials (Japanese and Taiwanese cultivars and Prunus relatives)

We used 112 Japanese and 5 Taiwanese cultivars of Japanese apricot (*P. mume*), and 7 apricot (*P. armeniaca*), 4 Japanese plum (*P. salicina*), and 1 peach (*P. persica*) cultivars (Table S1). For Japanese cultivars of *P. mume*, we used 55 fruit and 45 ornamental cultivars, 8 hybrids between *P. mume* and *P. armeniaca*, and 4 hybrids between *P. mume* and *P. salicina*. Cultivar categorization was based on previous reports (Hayashi *et al.*, 2008; Numaguchi *et al.*, 2019). All plant materials were maintained at the Japanese Apricot Laboratory, Wakayama Fruit Experiment Station (Minabe, Wakayama, Japan) as previously reported by Numaguchi *et al*. (2019) (Table S1).

### Target capture sequencing

Genomic DNA was extracted from young leaves using Nucleon PhytoPure (GE Healthcare, Chicago, IL, USA) and subjected to phenol/chloroform purification. We employed a KAPA HyperPlus kit (Kapa Biosystems, Wilmington, MA, USA) to construct gDNA-seq libraries for an Illumina platform. Libraries were barcoded for each sample using single 8-bp NEXTflex adaptors (Bioo Scientific, Austin, TX, USA) and enriched by PCR using PrimeSTAR Max (Takara Bio, Shiga, Japan) with the following protocol: 3 min at 95°C, followed by eight cycles of 10 s at 95°C, 30 s at 65°C, 30 s at 72°C, and final extension for 5 min at 72°C.

To selectively retrieve libraries with exons, a myBaits Custom design kit was used to design 1–20-K probes (Arbor Biosciences, Ann Arbor, MI, USA), which uses biotinylated RNA probes to concentrate fragments carrying sequences of interest (Gnirke *et al.*, 2009), based on the published genomic and coding sequences of *P. mume* (Zhang *et al.*, 2012). We selected 29,621 non-redundant coding loci showing single hits with BLAST+ (MEGABLAST with -p 70 option) against the *P. mume* genome, for the subsequent bait designing. A 120-mer bait with 25–55 GC% per locus was randomly designed for each locus, and finally we obtained a bait set targeting 15,171 coding loci. We pooled an equal amount of constructed Illumina libraries (eight samples per tube). Pooled libraries were purified by AMPure XP (Beckman Coulter, Indianapolis, IN, USA) and then electrophoresed on 1% agarose gel. We cut out a 300–700-bp area of DNA bands to re-extract libraries using a FastGene Gel/PCR extraction kit (NIPPON Genetics, Tokyo, Japan). Libraries were then subjected to target capture hybridization using myBaits Custom designed probes (Arbor Biosciences). Captured libraries were sequenced using the HiSeq 4000 platform (Illumina, San Diego, CA, USA) (paired-end 100 bp).

### SNP calling

In addition to our original sequencing data, we also used published sequencing data. Of the 348 *P. mume* cultivars in the whole-genome sequencing data reported by Zhang *et al*. (2018), 79 derived from China were selected and downloaded from Sequence Read Archive (https://www.ncbi.nlm.nih.gov/sra) (Table S2). The data were selected to evenly contain all the P1 to P16 phylogenetic clusters reported by Zhang *et al*. (2018). Importantly, the clade P1 contains interspecific hybrids such as *P. mume* × *P. armeniaca* and *P. mume* × *P. salicina*. Raw reads were trimmed with no-demultiplex-allprep-8 (https://github.com/Comai-Lab/allprep) to select the reads with high quality (Phred score > 20 over a 5-bp window, length > 35-bp) and containing no ‘N’ and adapter sequences. The selected reads were mapped against LG1–8 of the peach (*P. persica*) v2.0 reference genome (Verde *et al.*, 2017) using BWA-MEM with default parameters (Li, 2013). PCR duplicates were removed with OverAmp-3 (http://comailab.genomecenter.ucdavis.edu/index.php/Bwa-doall). SNPs were called using Samtools mpileup (Li *et al.*, 2009) and VarScan2 mpileup2snp (Koboldt *et al.*, 2009, 2012) with default settings (here, reffered to as “Primary_set”). From Primary_set, we removed loci with >20% missing genotyping rate with PLINK (Purcell *et al.*, 2007). Then, missing genotypes were imputed using Beagle 5.0 (Browning *et al.*, 2018) with default settings (“Imputed_set”). We also prepared “Cap_set” by removing the Chinese cultivars from Primary_set. Cap_set was subjected to sequencing depth estimation with VCFtools (Danecek *et al.*, 2011). Imputed_set and Cap_set were used to estimate annotated genomic locations using SnpEff (Cingolani *et al.*, 2012).

### Population structure analysis

SNP sets for population structure analyses were prepared based on Imputed_set. We first extracted cultivars of interest and removed loci with a minor allele frequency (MAF) <0.03 and that violated the Hardy–Weinberg equilibrium (*P* <0.0001) using PLINK. We further pruned SNPs with high (*r*^2^ >0.5) linkage disequilibrium (LD) within a 50-SNP window with 3 SNPs shifting using PLINK (--indep 50 3 2).

Population structure was estimated using three methods: principal component analysis (PCA), Bayesian clustering, and maximum likelihood phylogenetic analysis. PCA was performed using smartpca of EIGENSOFT (Patterson *et al.*, 2006). ADMIXTURE (Alexander *et al.*, 2009) was used for Bayesian clustering. We assumed *K* = 2–10, and 10 simulations were carried out for each *K* value. We then compiled the results of 10 simulations for each *K* using CLUMPP (Jakobsson & Rosenberg, 2007). The most optimal *K* was estimated based on cross-validation error (CVE) values calculated according to the ADMIXTURE manual. In the present study, the most optimal *K* was estimated to be four (CVE = 0.328). A maximum likelihood (ML) phylogenetic tree was constructed using SNPhylo (Lee *et al.*, 2014) with 1,000 bootstrap replications.

For detection of linkage disequilibrium, based on Imputed_set, we removed samples Chi_30, 202, 270, 283, 396, Jap_AM1–8, and SM1–4, which were previously considered to be interspecific hybrids (Hayashi *et al.*, 2008; Zhang *et al.*, 2018; Numaguchi *et al.*, 2019), and additionally Chi_250 and Jap_O2, which were newly classified as “Admixed” in the present study. Pairwise LD was computed using PopLDdecay (Zhang *et al.*, 2019) with -MaxDist 10000, -MAF 0.03, and -Het 0.75 options. The Plot_Multipop function was then used to calculate moving averages of LD for each 10-kb bin.

For detection of genetic differentiation and identity by descent (IBD), the same SNP set as that used for the above population structure analyses was used. Pairwise Weir and Cockerham weighted *F*_ST_ was calculated using VCFtools. IBD was estimated using pairwise pi-hat values from PLINK.

### Identification of selective sweeps

To estimate genomic regions that experienced natural or artificial selection, we used the following methods: composite likelihood ratio (CLR) (Nielsen *et al.*, 2005), nSL (Ferrer-Admetlla *et al.*, 2014), and XP-EHH (Sabeti *et al.*, 2007) tests. Of these, the CLR test is a method based on SFS, which detects deviations in allele frequency from neutrality at each site. This assumes that a selected allele is fixed in a population (as in annual crops). Conversely, nSL and XP-EHH analyses are based on extended haplotype homozygosity (EHH), which detects elongated linkage disequilibrium blocks around a selected core allele. EHH-based methods work well if a selected allele is not completely fixed in a population but is maintained in a heterozygous state (as in trees, humans, and other animals) (Sabeti *et al.*, 2007; Kosova *et al.*, 2010; Akagi *et al.*, 2016; Lee *et al.*, 2016; Nadachowska-Brzyska *et al.*, 2019). XP-EHH compares EHH values between paired populations (e.g., ancestral and derived populations) and can detect selective sweeps related to population differentiation.

SweeD (Pavlidis *et al.*, 2013) (with -grid 500 flag) was used for calculation of CLR values. Neutral thresholds were determined according to Nielsen *et al.* (2005). We first generated 1,000 simulated neutral genotype datasets using ms (Hudson, 2002), based on the observed number of polymorphic sites (S) and sample size (n) of each subpopulation. Next, we ran SweeD with -grid 500 flag using simulated genotype sets to obtain neutral CLR values. Neutral thresholds were determined as 99% percentile values for each subpopulation. We used selscan v 1.2.0a (Szpiech & Hernandez, 2014) to perform nSL and XP-EHH analyses. We assumed that genetic position was equal to physical position in the XP-EHH analysis. An SNP dataset was generated based on Imputed_set. We first extracted genotype sets for each subpopulation (China, Japan, Taiwan, ornamental, fruit, and small-fruit). Loci with MAF <0.03 were filtered for each dataset, and subsequently, haplotypes were phased using Beagle 5.0. Unstandardized nSL and XP-EHH values were Z-scored using the norm function of selscan and transformed to *P* values.

### Genetic differentiation in genome fragments among subgen. Prunus species

For SNP data preparation, we first removed admixed cultivars from Imputed_set and subsequently paired Chinese, Japanese, and Taiwanese cultivars with apricots (*P. armeniaca*) or Japanese plums (*P. salicina*). We then filtered loci with MAF <0.03 with PLINK. Here, we attempted to estimate introgressed genomic positions based on sliding window characterization for indices of genetic differentiation. To do this, we assessed the transition of three indices for population differentiation: (i) value of the first principal component (PC1) in PCA, (ii) Q value of the ADMIXTURE analysis with *K* = 2, and (iii) Jost’s *D* value (Jost, 2008), with the sliding window approach. We conducted “Bin-PCA” and “Bin-Admixture” analyses, which refer PCA from scikit-learn (Pedregosa *et al.*, 2011) and ADMIXTURE (Alexander *et al.*, 2009), to consecutively calculate PC1 and Q values, respectively. We used 1-Mb bin and 500-kb walking size in Bin-PCA and Bin-Admixture analyses. PC1 values were Z-transformed based on the equation: zPC1 = (PC1−μPC1)/σPC1. Here, μPC1 and σPC1 indicate the average and standard deviation of PC1, respectively. Q values of Bin-Admixture were transformed into absolute values of the difference between each Q value of *P. mume* individuals (Pm_indv.) and average Q values for *P. armeniaca* or *P. salicina* (related_ave.) as follows: dQ = |Q_Pm_indv._−Q_related_ave._|. For calculating Jost’s *D*, we used vcfWindowedFstats in pypgen 0.2.1 (https://pypi.org/project/pypgen/) with a window size of 1 Mb.

### Detailed analysis on the loci with selective sweep and interspecific introgression

To further examine the inferred region with characteristic selection or introgression, we focused on some specific regions. Especially, we narrowed the possibly introgressed regions on chromosome 8 with Bin-Admixture setting the 50-kb bin and 25-kb walking size. We removed bins with <10 SNPs. Neighbor-joining phylogenetic analysis was performed to visualize the allelic evolution in the specific regions with TASSEL 5.0 (Bradbury *et al.*, 2007), using extracted SNPs of interested regions from Imputed_set. Roots for phylogenetic trees were determined at midpoints using FigTree v1.4.4 (https://github.com/rambaut/figtree/releases).

## Results

### Efficacy of targeted resequencing in the subgenus Prunus

From the targeted resequencing of 129 *Prunus* cultivars (117 *P. mume*, 7 *P. armeniaca*, 4 *P. salicina*, and 1 *P. persica*), 1,096,007,397 of a total 1,177,780,940 reads (93.1%) (deposited at DRA009691; Table S1) were mapped onto LG1–8 of the peach v2.0 genome, including 402,859,421 uniquely mapped reads (34.2%). In Cap_set, we could identify a total of 489,420 SNPs with an average depth of 29.8×, of which each cultivar ranged from 15.2× to 60.2×. SnpEff analysis revealed that total 94.4% of SNPs were located on a genic region (35.3%) or its upstream and downstream regions (26.8% and 32.3%, respectively) (Table S3). In Imputed_set, we obtained a total of 148,953 SNPs, of which the SnpEff result was almost consistent with that of Cap_set (Table S3). Thus, we could successfully and cost-effectively obtain SNPs based on exon capture in subgen. *Prunus* cultivars.

### Definition of population structure

A total of 14,310 selected SNPs were used for PCA, ADMIXTURE and ML phylogenetic analyses to reveal the population structure among 208 *Prunus* cultivars (79 Chinese cultivars were added to the 129 cultivars). In all three analyses, Japanese apricot (*P. mume*), apricot (*P. armeniaca*), and Japanese plum (*P. salicina*) were clearly found to form species specific clusters (Figs. 1, 2). This was also supported by the *F*_ST_ values among the species (Table S4). Hypothetical interspecific hybrids of *P. mume* (Jap_AM1–8, SM1–4, Chi_30, 202, 270, and 396) (Hayashi *et al.*, 2008; Zhang *et al.*, 2018; Numaguchi *et al.*, 2019) were positioned between *P. mume* and the other *Prunus* species, supporting that they are “Admixed” individuals (Figs **1**, **2**). Importantly, Chinese and Japanese cultivars of *P. mume* were clustered into separate groups, whereas Taiwanese cultivars were clustered with Japanese cultivars (Figs **1a, b**, S1). The ML tree (Fig. **2**) also supported that the *P. mume* cluster was largely divided into Chinese and Japanese (with Taiwanese) clades with statistical support (bootstrap >60 in ML), including minor exceptions (Jap_O6, 9, 14, 18, 27, 42, 44, and 45; green stars in Fig. **2**). In the IBD analysis, some pairs of Japanese and Taiwanese cultivars were inferred to be in first- or second-degree relationships (pi-hat = 0.25–0.5; Fig. S2). Conversely, some pairs of Chinese and Japanese cultivars showed first-degree relationships (pi-hat = 0.5), but most combinations were genetically distinct (Fig. S2). The highest *F*_ST_ value was observed between Chinese and Taiwanese cultivar groups, in contrast to the geographical proximity (Table S4). These results differ from conventional (or empirical) observations, which have indicated that Japanese cultivars of *P. mume* were originally introduced from China to Japan relatively recently (ca. 2,000 years ago) via human activities (Horiuchi *et al.*, 1996). Based on the ML tree, Japanese cultivars showed weak differentiation depending on their characteristics or application by humans (Fig. **2**). Although subpopulations of fruit, small-fruit, and ornamental cultivars was not clearly divided in PCA, ADMIXTURE, and *F*_ST_ (Figs S1, **1b**, Table S5), in the ML tree, the majority of fruit (36 of 45 cultivars), small-fruit (9 of 10 cultivars), and ornamental (25 of 45 cultivars) cultivars belonged to the same cluster (Fig. **2**). This supports the possibility that human preference triggered a recent differentiation of Japanese population from the same genetic resources (Horiuchi *et al.*, 1996; Hayashi *et al.*, 2008; Numaguchi *et al.*, 2019).

**Fig. 1.**
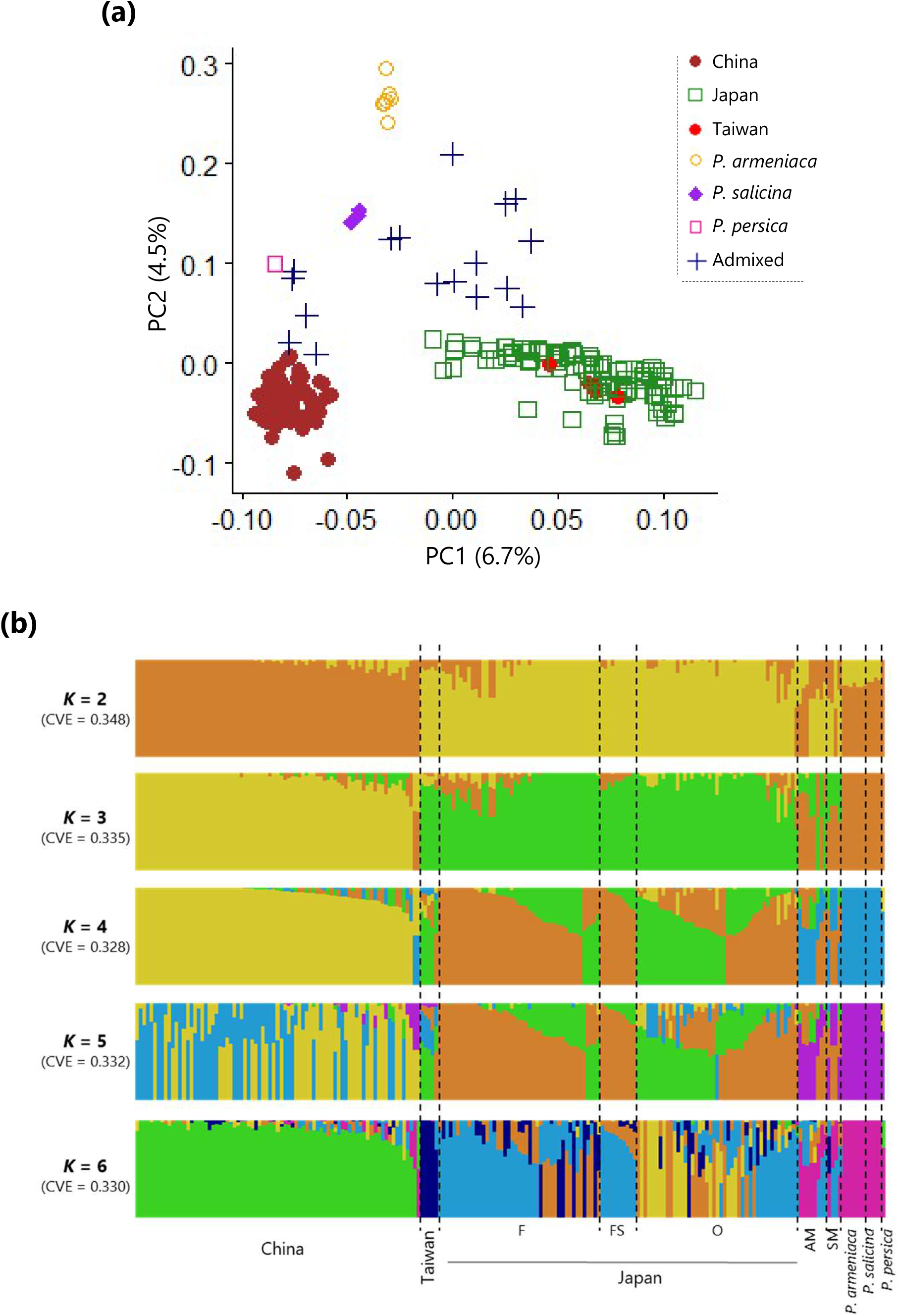
Population structure analysis in *Prunus mume*. (**a**) Principal component analysis (PCA) of all the 208 *Prunus* cultivars. (**b**) Proportion of ancestry for all the 208 *Prunus* cultivars from *K* = 2–6 inferred with ADMIXTURE. Proportions of the membership to each cluster are shown with the lengths of the colored bar (y-axis). CVE: cross validation error. F: fruit cultivars, FS: small-fruit cultivars, O: ornamental cultivars, AM: putative hybrids with *P. armeniaca*, SM: putative hybrids with *P. salicina*.

**Fig. 2.**
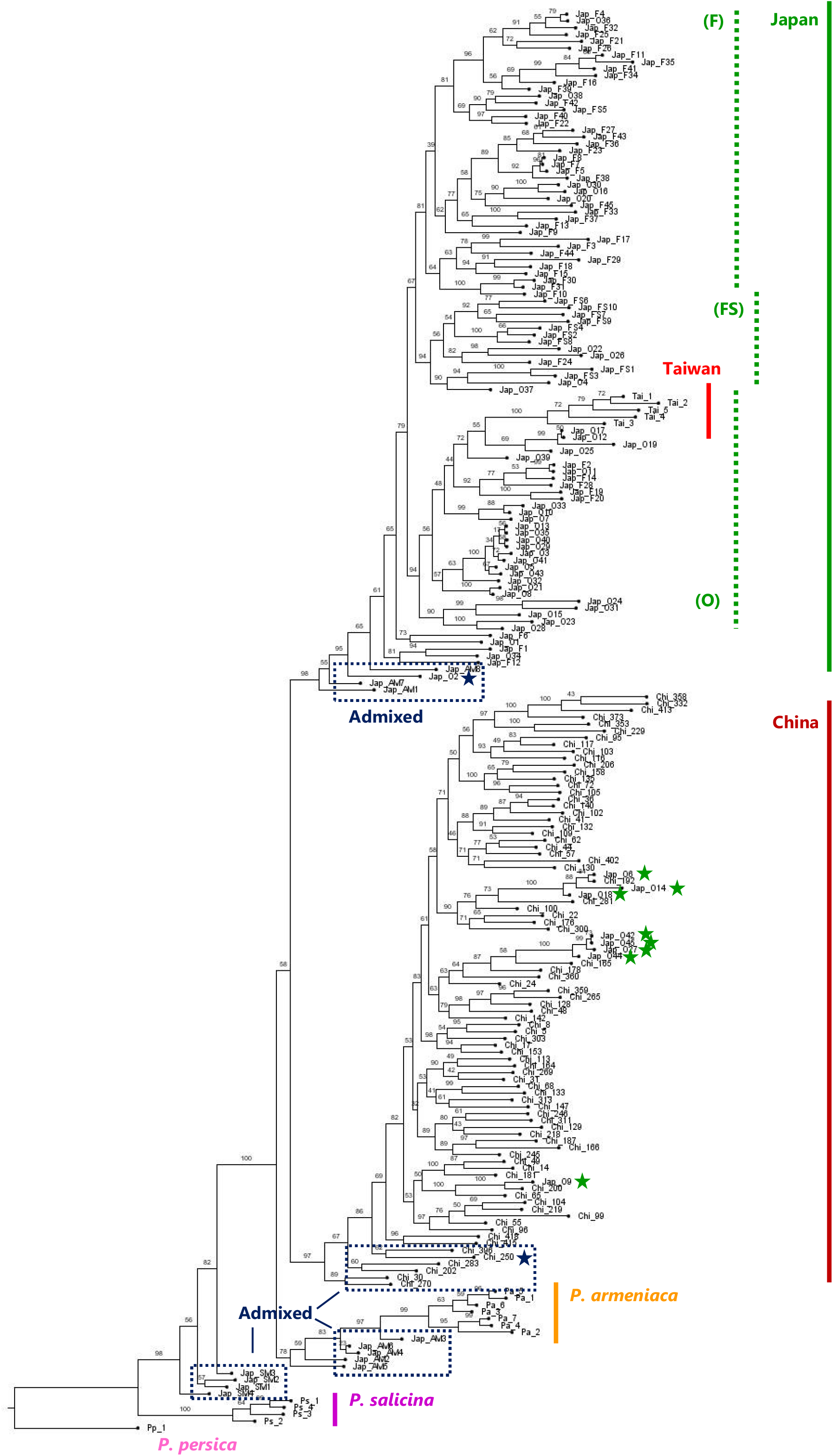
Maximum likelihood phylogenetic tree inferred with all the 208 *Prunus* cultivars. Values beside the nodes indicate bootstrap values generated with 1,000 replications. *P. mume* cultivars clustered with admixed cultivars are shown with navy stars. Dotted lines within Japanese clusters indicate the characteristic clusters for fruit (F), small-fruit (FS), and ornamental (O) cultivars. Green stars indicate Japanese cultivars in Chinese clusters. Chi: Chinese cultivars, Jap: Japanese cultivars, Tai: Taiwanese cultivars, F: fruit cultivars, FS: small-fruit cultivars, O: ornamental cultivars, AM: putative hybrids with *P. armeniaca*, SM: putative hybrids with *P. salicina*, Pa: *P. armeniaca*, Ps: *P. salicina* and Pp: *P. persica*.

LD mostly decayed within ca. 100 kb in all the *P. mume* groups surveyed in the present study (Figs **3**, S3), which is much longer than in *P. armeniaca* but shorter than in *P. persica* (Mariette *et al.*, 2016; Akagi *et al.*, 2016; Yu *et al.*, 2018). The extent of LD was slightly different among cultivar groups. For example, LD decayed slower in Japanese cultivars than in the others (Fig. **3**). Within the Japanese cultivars, ornamental cultivars exhibited further slower LD decay (Fig. S3), presumably due to their narrow genetic resources and frequent utilization of bud-sport for development of new cultivars especially after the Edo Period (Mega *et al.*, 1988; Horiuchi *et al.*, 1996).

**Fig. 3.**
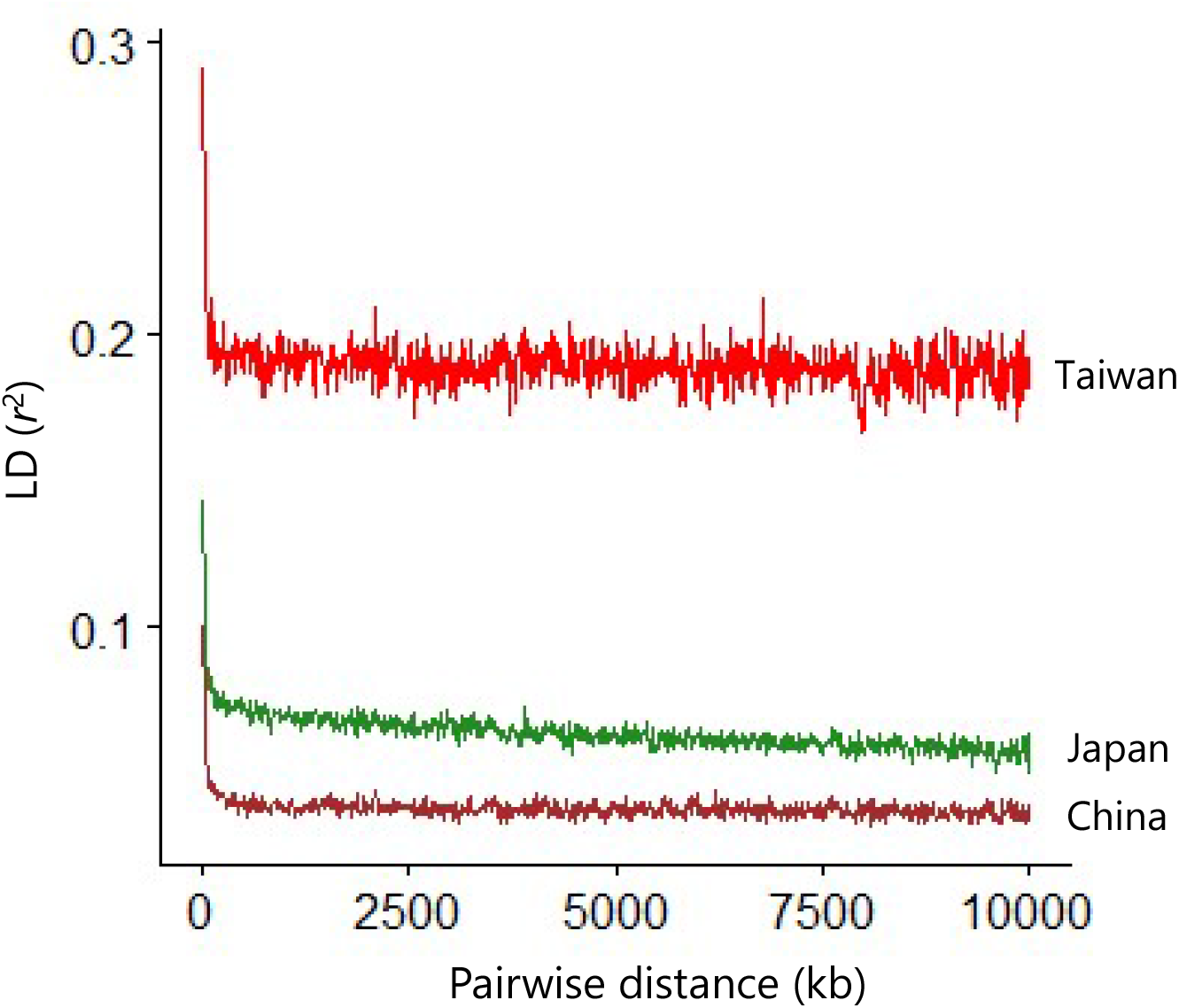
Patterns of linkage disequilibrium decay in Chinese, Japanese, and Taiwanese cultivars of *Prunus mume*.

### Identification of selective sweeps related to population differentiation in P. mume

Alleles that have undergone positive selection showed, i) reduction in genetic diversity, and ii) extension of haploblock, in adjacent genetic regions and a targeted locus itself (Stephan, 2019). We applied an SFS-based method, SweeD, to detect i), whereas we used extended haplotype homozygosity (EHH)-based method, nSL and XP-EHH, to identify ii).

For Chinese and Japanese populations, the results for SweeD and single-population EHH-based nSL analyses were different (Figs **4a**, S4, Table S6). Especially in the Chinese population, no SweeD peak exceeded the neutral threshold (Fig. S4). Tree crops have specific characters, such as long generation time and frequent vegetative propagation, suggesting that selected alleles may have not been completely fixed, like in humans (Voight *et al.*, 2006; Akagi *et al.*, 2016). Sites with only SFS-based peaks (with no EHH-based peaks) are thus suspected to be derived from the occasional reduction of nucleotide diversity, namely, the substitution ratio in the *P. mume* genome, distortion of the availability in SNPs, or simple drift (Akagi *et al.*, 2016). Therefore, in *P. mume*, we mainly focused on the results of the EHH-based analyses in the following sections. Only in the Taiwanese cultivar group was the pattern of significant peaks of SweeD similar to that of nSL (Figs **4a**, S4, Table S6), suggesting that Taiwanese cultivars may have experienced stronger selection pressure than the other cultivar groups.

**Fig. 4.**
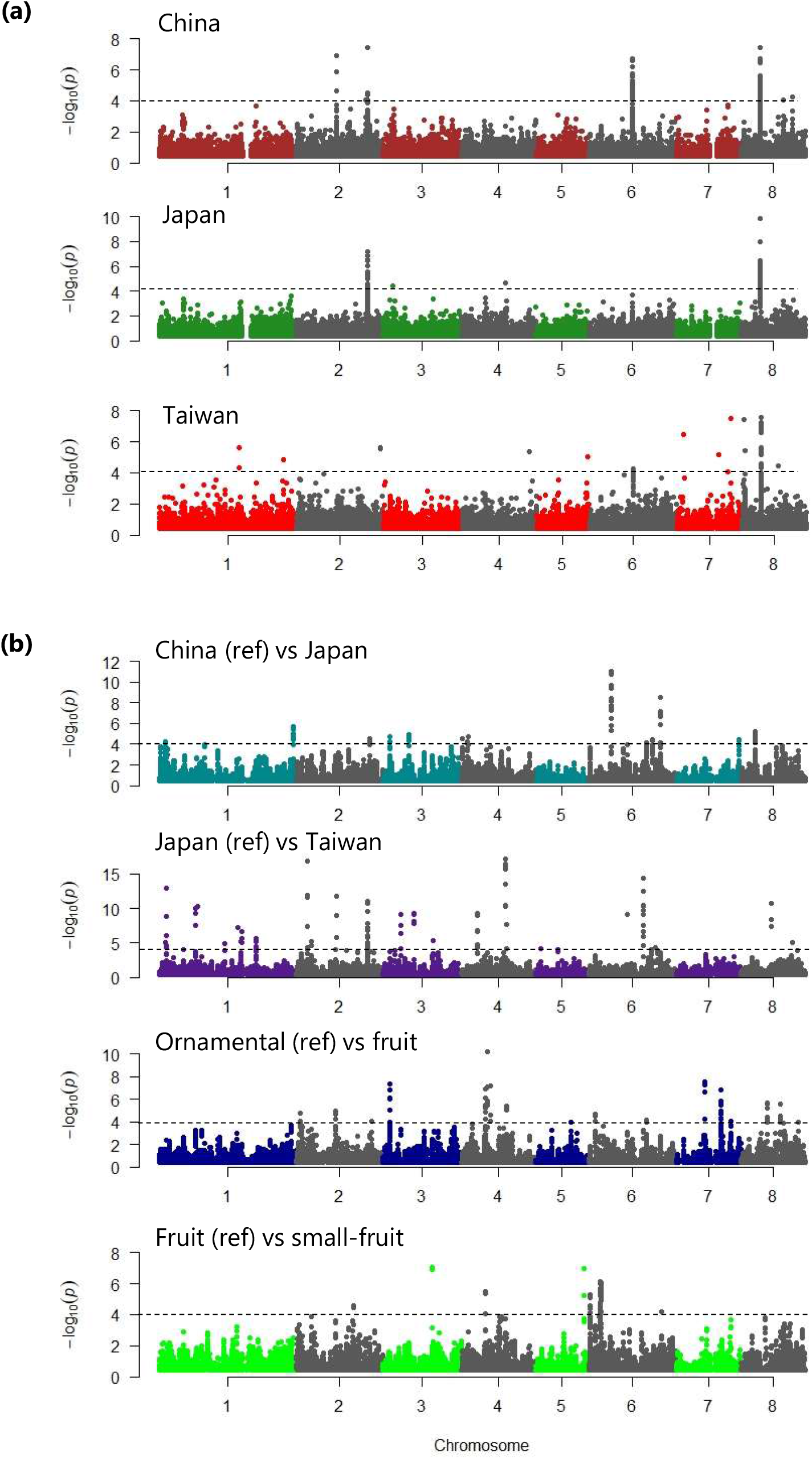
Identification of selective sweeps in Chinese, Japanese, and Taiwanese cultivars of *Prunus mume* based on, (**a**) single-population nSL and (**b**) dual-population XP-EHH analyses. *P* values calculated with normalized nSL values are shown in −log_10_ scale. Detailed information of SNPs with −log_10_*P* > 4 (dotted line) is summarized in Tables S6 (nSL) and S7 (XP-EHH).

In the nSL analysis, we could identify a strong peak (*P* < 1e^−4^) on chromosome 8 (ca. 6.7–6.8 Mb), which was common in all Chinese, Japanese, and Taiwanese cultivar groups of *P. mume* (Fig. **4a**). A peak on chromosome 6 was also common in three geographic groups, but the Japanese peak was not significant. Strong peaks common in Chinese and Japanese cultivars were observed on chromosome 2. Geographically specific peaks were found on chromosomes 2 and 8 in Chinese, chromosomes 3 and 4 in Japanese, and chromosomes 1, 2, 4, 5, 7, and 8 in Taiwanese populations. These results suggest that the common ancestor of *P. mume* underwent certain positive selection, such as on chromosomes 6 and 8, and thereafter established the three populations based on geographical separation and independent selection.

Furthermore, we conducted two population-based XP-EHH analyses in, (i) the Chinese cultivars (reference) vs. the Japanese cultivars (derived), (ii) the Japanese cultivars (reference) vs. the Taiwanese cultivars (derived), (iii) the ornamental cultivars (reference) vs. fruit cultivars (derived), and (iv) the fruit cultivars (reference) vs. the small-fruit cultivars (derived) (Fig. **4b**). The XP-EHH can focus on the selected alleles that are highly differentiated between populations (Sabeti *et al.*, 2007). Here, negative and positive normalized XP-EHH values indicated that an extended haploblock was observed in the reference and derived populations, respectively (Szpiech & Hernandez, 2014). Most of the strong peaks (*P* < 1e^−4^; Table S7) were not overlapped with those detected in the nSL analysis (Fig. **4b**). In the Chinese vs Japanese analysis, significant peaks were observed on all chromosomes except for chromosome 5 (Fig. **4b**). The genomic positions of these peaks were different from those in SweeD (Fig. S4, Table S7), indicating that they have not yet perfectly fixed in the population. Most strong XP-EHH peaks in Japanese vs Taiwanese groups showed negative values, except for ca. 33.9 Mb of chromosome 1 (Table S7), indicating that Japanese cultivars underwent much more extensive selection in the differentiation from Taiwanese cultivars. In the analyses of Japanese ornamental vs fruit cultivars and fruit vs small-fruit, we could also find many significant peaks, presumably involving ongoing selection in favor of human preference in Japan. Especially, peaks of ca. 3.9 Mb in chromosome 6 overlapped with the SweeD peak (Figs **4b**, S5, Table S7), suggesting strong selection pressure for the small-fruit trait in Japanese cultivars. Accordingly, tests for selection in *P. mume* populations identified its tree-crop-specific patterns for natural or artificial selection.

### Fragmental interspecific introgression in the subgen. Prunus

We detected, in 1-Mb bins, the zPC1 (Bin-PCA) and proportion of the Q values in Admixture with *K* = 2 (Bin-Admixture) to scan for genome-wide transition of genetic differentiation or potential introgressions for *P. mume* vs *P. armeniaca* or *P. salicina*. The Bin-PCA and Bin-Admixture showed mostly consistent results (Figs S6, S7). In most of the chromosomes, *P. mume* genomes showed signs of fragmental interspecific introgression (or no clear differentiation between species) from *P. armeniaca* (Fig. S6) or *P. salicina* (Fig. S7). When the three geographic groups were compared, Japanese cultivars showed the most frequent signals of interspecific introgressions from *P. armeniaca* or *P. salicina* (Figs S6b, S7b), while Taiwanese cultivars rarely showed them (Figs S6c, S7c). We also calculated a distance-matrix-based Jost’s *D* statistic in 1-Mb bins. Jost’s *D* values tended to be low in the genomic regions with interspecific introgression signals in Bin-PCA and Bin-Admixture (Figs S6, S7). However, unlike Bin-PCA and Bin-Admixture, the transition of Jost’s *D* values substantially fluctuated in most chromosomes (Figs S6, S7). The overlapped region of Bin-PCA, Bin-Admixture, and Jost’s *D* signals may be a strong signature of interspecific introgressions, indicating the especially low allelic divergence between *P. mume* and relatives.

Importantly, some signals of interspecific introgressions were overlapped with the selective sweep (nSL peaks) (Fig. **5**; hereafter, “introgression-sweep”), indicating that introgressed regions may have been positively selected in the evolution of *P. mume* populations. They were located on chromosomes 6 and 8 in Chinese cultivars, 2, 3, 4, 6, and 8 in Japanese cultivars, and 6 and 8 in Taiwanese cultivars. In chromosomes 6 and 8, nSL signals harbor fragment introgressions from both *P. armeniaca* and *P. salicina* (Fig. **5a-c**). In chromosome 2, 3, and 4 of Japanese cultivars, introgression signals were accompanied by nSL peaks independently of other groups (Fig. **5b**). These results suggest that interspecific introgressions independently contributed to the establishment of not only *P. mume* but also of each geographical group following the divergence of these subpopulations.

**Fig. 5.**
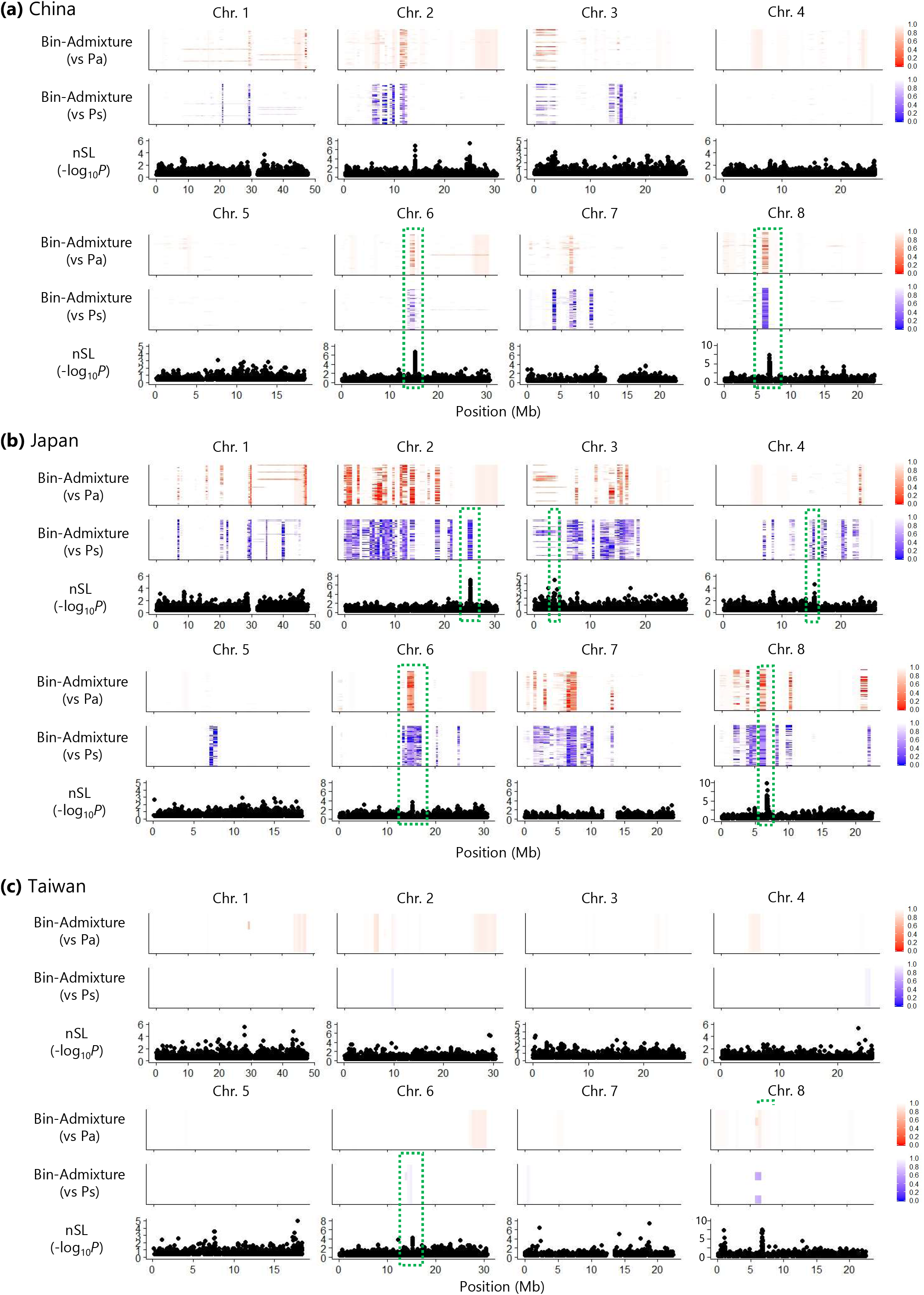
Examination of conformity between putative interspecific introgression and selective sweep signals. Genomic locations of Bin-Admixture signals were compared with those of nSL peaks. Bin-Admixture (1-Mb-binned) analyses with *Prunus armeniaca* (Pa, red signals) and *P. salicina* (Ps, blue signals), and nSL scans were performed in, (**a**) Chinese, (**b**) Japanese, and (**c**) Taiwanese cultivars of *P. mume*. The degree of introgression is indicated by the color scales to the right (0: highly introgressed–1: not introgressed). Potential introgression-sweep regions are highlighted by green dotted rectangles.

It is worth noting that the strongest introgression-sweep signal was detected around 6 Mb on chromosome 8 (Fig. **6a,b**). Fine assessment of the Q values with shorter bins (50 kb) increased the resolution for the genomic region with overlap of Bin-Admixture and nSL signals around 6.7–7.1 Mb (Fig. **6c**). Next, we compared evolutionary topologies constructed from the SNPs in region 1 (6.0–6.7 Mb), region 2 (6.7–7.1 Mb), region 3 (7.1–8.5 Mb), and in the whole chromosome 8, according to an approach by Choi & Purugganan (2018) (Fig. **6d–g**). The topology for the whole chromosome 8 (Fig. **6d**) was mostly consistent with that for the whole genome (Fig. **2**) in accordance with the divergence of species and populations. For regions 1–3, only region 2 showed a distinct topology with alleles of *P. mume* and related species in the subgenus *Prunus* grouped together, while alleles putatively introgressed and under selection were nested in a single clade with *P. salicina* alleles (Fig. **6f**), showing very small genetic differentiation (alleles with green band in Fig. **6f**). Consequently, region 2 (6.7–7.1 Mb) is thought to have been exposed to positive selection pressure, which may be associated with the adaptive evolution of *P. mume* to import advantageous alleles from other species.

**Fig. 6.**
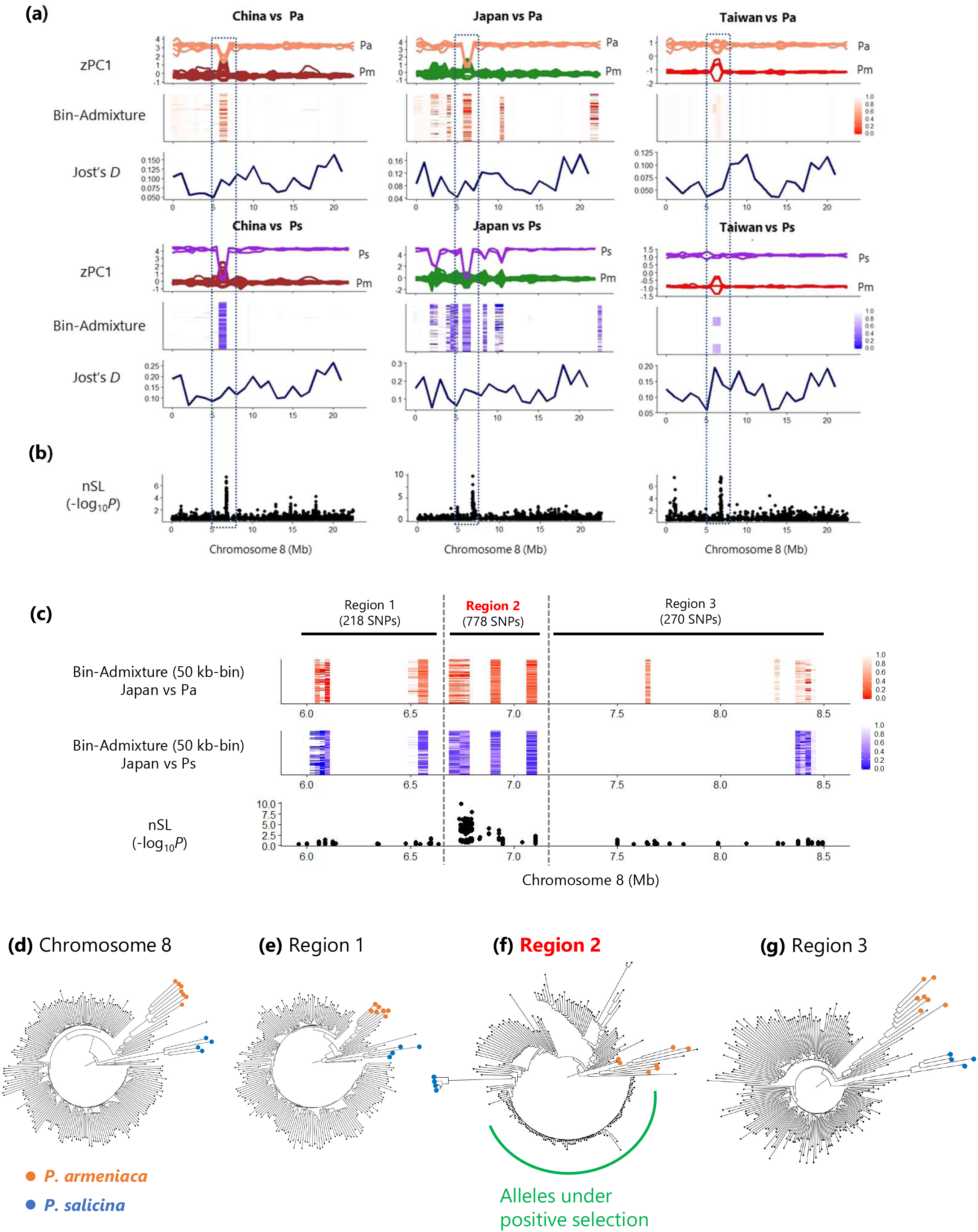
Consistent region for interspecific introgression and selective sweep on chromosome 8. (**a**) Bin-PCA, Bin-Admixture, and Jost’s *D* patterns in Chinese, Japanese, and Taiwanese cultivars, in comparison to *Prunus armeniaca* (Pa, upper panels) and *P. salicina* (Ps, lower panels). The degree of introgression in Bin-Admixture is indicated by the color scales to the right (0: highly introgressed–1: not introgressed). (**b**) Transitions of nSL for an index of selective sweep. *P* values calculated with normalized nSL values were shown in −log_10_ scale. (**c**) Close up of the genomic fragment (ca. 6.0–8.5Mb) showing overlapped nSL peaks and potential interspecific-introgression in Japanese cultivars of *P. mume*. We further divided this region into three sub-fragments (Regions 1–3) according to the pattern of nSL plots to assess their phylogenetic relationships. (**d**) Neighbor-joining phylogenetic trees with the whole SNPs in chromosome 8 and with the SNPs in Regions 1–3. The tree for Region 2 showed a topology inconsistent with the whole chromosome 8 and the flanking regions (Regions 1 and 3), and was also inconsistent with the estimated speciation pattern of the subgenus *Prunus*. Alleles that underwent potential selective sweeps were indicated with a green solid line.

## Discussion

### Completed and ongoing population differentiation in P. mume

In the present study, we revealed that Chinese and Japanese cultivars of *P. mume* showed distinct population differentiation (Figs **1**, **2**). Conversely, Taiwanese cultivars belonged to the Japanese clade but clustered independently from the Japanese cultivars, consistent with the results of previous studies (Hayashi *et al.*, 2008; Numaguchi *et al.*, 2019). These results suggest that the differentiation of Chinese and Japanese populations predated that of the Taiwanese population, which is inconsistent with the conventional belief that *P. mume* cultivars were derived from Chinese ones, and recently (ca. 2000 years ago), people have been introduced to other regions (Mega *et al.*, 1988; Faust *et al.*, 2011). There is a record describing wild *P. mume* accessions collected from western Japan that have been clonally maintained (https://agriknowledge.affrc.go.jp/RN/3030041889; in Japanese). Those samples may have important genetic information related to the origin of Japanese cultivars. We may directly investigate the population structure of domesticated crops using ancient DNA (aDNA) extracted from ancient plant remains (Frantz *et al.*, 2016; Mascher *et al.*, 2016; Kistler *et al.*, 2018; Narasimhan *et al.*, 2019; Allaby *et al.*, 2019; Smith *et al.*, 2019). Thus, collaboration between geneticists and archaeologists will make rapid progress in studies on crop domestication.

### Contribution of genomic fragments undergoing interspecific introgressions and positive selections in the evolution of P. mume

We observed a higher number of strong (and successive) peaks in EHH-based scans than in an SFS-based SweeD analysis. This suggested that, in tree (or perennial) crops, most selected alleles are maintained in a heterozygous state, as suggested previously by Akagi *et al*. (2016). Since most SNPs called in our exon capture approach were located around protein coding regions (Table S3), it is expected that we may be able to detect functional nucleotide polymorphisms (FNPs) even with low sequencing coverage for each individual (Bamshad *et al.*, 2011; Kaur & Gaikwad, 2017). We could detect candidate genes potentially involved in environmental adaptation (Fig. **4**, Tables S6, S7). For instance, in the regions with strong selective sweep, leucine rich repeat containing proteins (e.g., Prupe1G161800, Prupe4G157900, Prupe8G046600, Prupe.8G012000, and Prupe.8G012800) and receptor-like kinases (e.g., Prupe.6G183600 and Prupe.6G261400) would commonly contribute to pathogen recognition pathways (Ellis *et al.*, 2000). The BTB/POZ-MATH-TRAF-like protein (Prupe.1G107200) is potentially associated with virus resistance in *P. armeniaca* (Mariette *et al.*, 2016). Genes potentially involved in stress response (e.g., Prupe.2G089100, Prupe.2G145200, and Prupe.3G110300) (Vij & Tyagi, 2008; Cheng *et al.*, 2011) were also identified (Tables S6, S7). These results suggest that selection on biotic or abiotic stress responsive genes may have contributed to the geographic separation of Chinese, Japanese, and Taiwanese cultivars.

According to the results of Bin-PCA, Bin-Admixture, and Jost’s *D* analyses, it was suggested that substantial fractions of *P. mume* genomes have frequently exchanged genomic fractions with related species of the subgen. *Prunus*. Large fractions of introgression may indicate that *P. mume*, especially in Japanese cultivars, have experienced a limited number of generations since the interspecific hybridization. Two genomic regions on chromosomes 6 and 8 were detected to have interspecific introgressions in Chinese, Japanese, and Taiwanese cultivar groups (Figs 5, 6). Particularly, a 6.74–6.80-Mb region in the region 2 on chromosome 8 contained high nSL peaks commonly detected among three groups (Table S6). Although this region carries no genes in the reference peach (*P. persica*) genome, we can propose the two following possibilities: 1) only in the *P. mume* genome, the selected haploblock in the corresponded region harbors candidate genes, and 2) this region includes *cis*-regulatory elements affecting gene expression in the flanking regions. For hypothesis 2), often, *cis*-elements were located distantly upstream (>10 kb) of the genes (Clark *et al.*, 2004; Konishi *et al.*, 2006; Ishii *et al.*, 2013; Ricci *et al.*, 2019). Prupe.8G057100 (mitochondrial transcription termination factor) is located ca. 80 kb from the selected haploblock (Fig. S8). Mitochondrial transcription termination factors have been reported to be related to abiotic stress response by controlling the expression level of nuclear genes (Quesada, 2016). Another region on chromosome 6 (ca. 15.2–15.3 Mb) also showed common introgression-sweep (Figs **5**, S9, Table S6). Although this region also harbored no annotated genes in the peach reference genome as well as the described introgression-sweep region on chromosome 8, it may have been important in the evolution of *P. mume*.

Other than the introgression-sweep commonly underwent among the geographical cultivar groups, our exon capture sequencing would allow efficient identification of FNPs in introgression-sweep regions specific to each cultivar group. Japanese cultivars have the largest fractions of interspecific introgressions, and they also show several geographically unique introgression-sweep regions in nSL (Fig. **5b**, Table S6). Of them, interleukin-1 receptor-associated kinase 1 (IRAK1) (Prupe.2G223200), which harbors the introgression signal of *P. salicina* on chromosome 2 (ca. 25 Mb), may act for pathogen recognition pathways (Jebanathirajah *et al.*, 2002; Dardick & Ronald, 2006). On chromosome 3 (ca. 3.6 Mb), 3-epi-6-deoxocathasterone 23-monooxygenase (Prupe.3G050900) was reported to be associated with brassinosteroid biosynthesis (Ohnishi *et al.*, 2006) and is potentially involved in the fruit-enlargement process. Premnaspirodiene oxygenase (Prupe.4G237900) on chromosome 4 (ca. 15.5 Mb) is a kind of cytochrome P450, which participates in terpene biosynthesis (Weitzel & Simonsen, 2015). Terpenes are the largest class of plant-derived compounds that have numerous potential applications across food, beverage, pharmaceutical, cosmetic, and agriculture industries (Boutanaev *et al.*, 2015). These results suggest that Japanese cultivars might import genetic factors from other species to satisfy the preference of people in Japan.

### Inferring the evolution of P. mume

The results so far propose an evolutionary model for the establishment of the current *P. mume* populations, where frequent interspecific introgressions with natural/artificial selection have played important roles (Fig. **7**). Several hybridization events might have occurred in the differentiation of these three species of the subgenus *Prunus* (*P. mume*, *P. armeniaca*, and *P. salicina*). During the formation of the *P. mume* common ancestor, important introgressions related to unknown phenotypic changes on chromosomes 6 and 8 (Figs **5**, **6**, S9) were positively selected. After the establishment of the original *P. mume*, three core populations were further differentiated to be adapted to China, Japan, and Taiwan, in which independent introgressions from *P. armeniaca* or *P. salicina* and positive selection on some regions might have contributed to the establishment of each population (e.g., chromosomes 2, 3, and 4 in Japanese cultivars, Fig. **5b**). Recently, the *P. mume* Japanese cultivars differentiated to form sub-populations, such as ornamental, fruit, and small-fruit clusters, based on human-preference-associated selection (Fig. **4b**, Table S7).

**Fig. 7.**
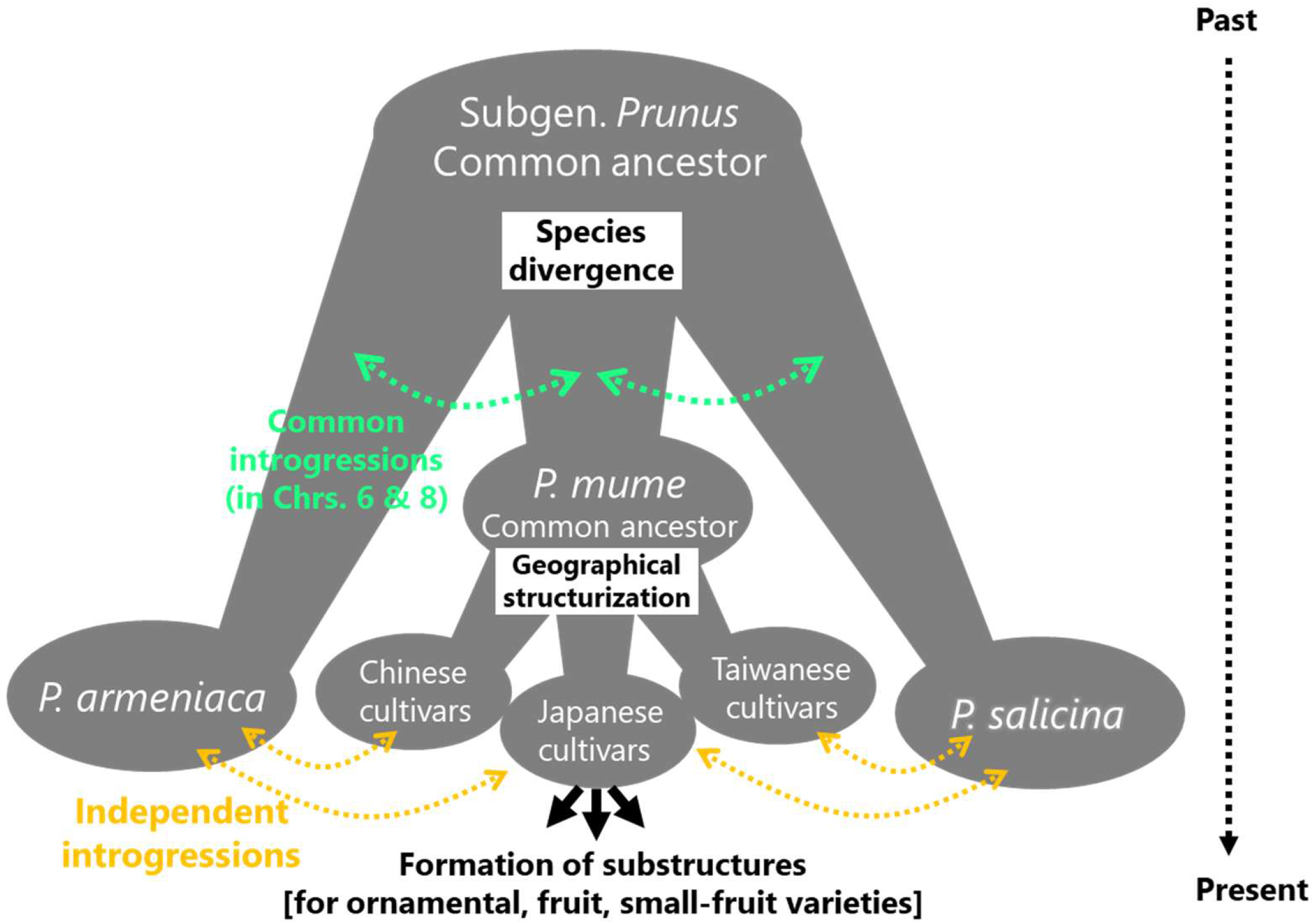
Tentative process of evolution in *Prunus mume*. The three subgenus *Prunus* species, Japanese apricot (*P. mume*), apricot (*P. armeniaca*), and Japanese plum (*P. salicina*), may have diverged from their common ancestor, experiencing mutual hybridizations. Important introgressions commonly detected on chromosomes 6 and 8 may have been selected during the formation of the *P. mume* common ancestor. *P. mume* may have further experienced independent introgression and selection, resulting in differentiation based on geographical separation and human preference.

The emphasis of the present study was the finding that there are many naturally/artificially selected regions derived from interspecific introgressions, and that they have considerably contributed to the establishment of current *P. mume* populations. Interspecific introgressions are thought to also be important in the evolution of other annual or perennial crops, such as rice (Choi & Purugganan, 2018), wheat (He *et al.*, 2019), maize (Hufford *et al.*, 2013; Brandenburg *et al.*, 2017), apple (Cornille *et al.*, 2012), and olive (Diez *et al.*, 2015; Gros-Balthazard *et al.*, 2019). These reports pointed out the importance of crop-wild introgressions to transfer wild beneficial alleles into domesticates and vice versa, which enabled them to rapidly expand and adapt to new climatic and agricultural conditions. Notwithstanding, especially in woody crops, the genomic landscape of interspecific introgressions and their actual contribution to evolution remains poorly understood. Our findings shed light on the complicated nature of adaptive evolution with interspecific introgressions among domesticated tree crops.

## Acknowledgements

We are deeply grateful to researchers and technicians at the Japanese Apricot Laboratory, Wakayama Fruit Tree Experiment Station for maintaining the *Prunus* cultivars. We used the Vincent J. Coates Genomics Sequencing Laboratory at UC Berkeley, supported by NIH S10 OD018174 Instrumentation Grant. This work was supported by JSPS KAKENHI Grant Number JP18K14449 to K. N. The English language of this paper was edited by Editage (https://www.editage.com/).

## Author contributions

K. N. and T. A. planned and designed the research. K. N., Y. K. and R. I. carried out the targeted resequencing experiments. K. N. analyzed the data. T. A. developed bioinformatic approaches. K. N., T. A. and T. I. wrote and revised the manuscript.

## Data availability statement

Raw fastq reads for 129 *Prunus* cultivars resequenced in this study were deposited in the Sequence Read Archive (SRA) under DRA accession number DRA009691 (Bioproject accession number PRJDB9365). Sources for all downloaded data are referred to in the Supplementary information.

## Supporting Information

Additional supporting information may be found in the online version of this article.

**Fig. S1** Principal component analysis (PCA) of Japanese and Taiwanese cultivars of *Prunus mume*.

**Fig. S2** Pairwise identity by descent (IBD) proportions in *Prunus mume* cultivars.

**Fig. S3** Patterns of linkage disequilibrium decay among Japanese cultivars of *Prunus mume*: fruit (F), small-fruit (FS) and ornamental (O) cultivars.

**Fig. S4** Identification of selective sweeps in Chinese, Japanese, and Taiwanese cultivars of *Prunus mume* based on site frequency spectrum (SFS)-based SweeD (composite likelihood ratio, CLR) analysis.

**Fig. S5** Identification of selective sweeps in fruit, small-fruit, and ornamental cultivars of Japanese *Prunus mume* based on site frequency spectrum (SFS)-based SweeD (composite likelihood ratio, CLR) analysis.

**Fig. S6** Chromosomal patterns of genetic differentiation among, (a) Chinese, (b) Japanese, and (c) Taiwanese cultivars of *Prunus mume* and *P. armeniaca*.

**Fig. S7** Chromosomal patterns of genetic differentiation among, (a) Chinese, (b) Japanese, and (c) Taiwanese cultivars of *Prunus mume* and *P. salicina*.

**Fig. S8** Positions of annotated genes adjacent to the candidate region (red) on chromosome 8.

**Fig. S9** Neighbor-joining phylogenetic trees with the single nucleotide polymorphisms (SNPs) in (a) 15.2–15.3, in its (b) upstream and (c) downstream 1-Mb regions and (d) with the whole SNPs in chromosome 6.

**Table S1** Japanese and Taiwanese cultivars and other *Prunus* species used in the present study.

**Table S2** Chinese cultivars (after Zhang *et al*., 2018) used in the present study.

**Table S3** Percentage of genomic locations of single-nucleotide polymorphisms (SNPs) derived from targeted resequencing.

**Table S4** Pairwise *F*_ST_ among *Prunus* cultivars.

**Table S5** Pairwise *F*_*ST*_ among Japanese *Prunus mume* cultivars.

**Table S6** The strongest candidates for selective sweep based on the nSL test for selection.

**Table S7** The strongest candidates for selective sweep based on the XP-EHH test for selection.

