## Supplementary Figures for "Interspecific introgression and natural selection in the evolution of Japanese apricot (*Prunus mume*)"

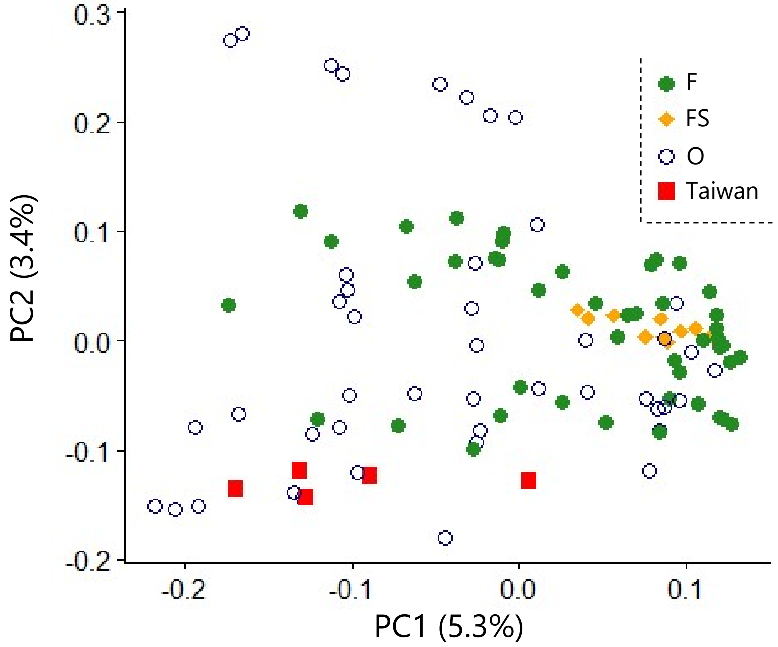


**Fig. S1** Principal component analysis (PCA) of Japanese and Taiwanese cultivars of *Prunus mume*. Percentage of the total variation explained by the PC is shown in parentheses. F: fruit cultivars, FS: small-fruit cultivars, O: ornamental cultivars.

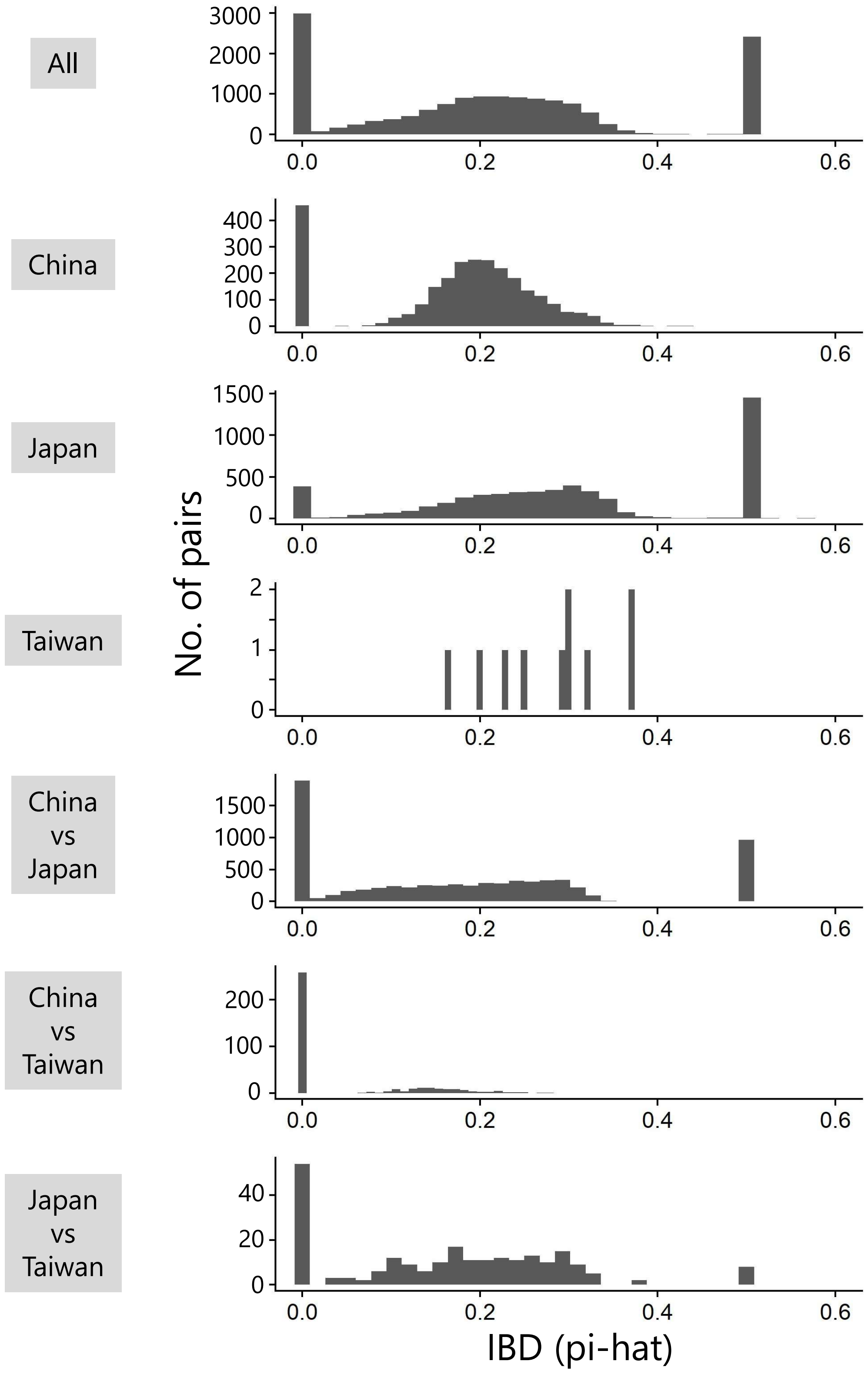


**Fig. S2** Pairwise identity by descent (IBD) proportions in *Prunus mume* cultivars.

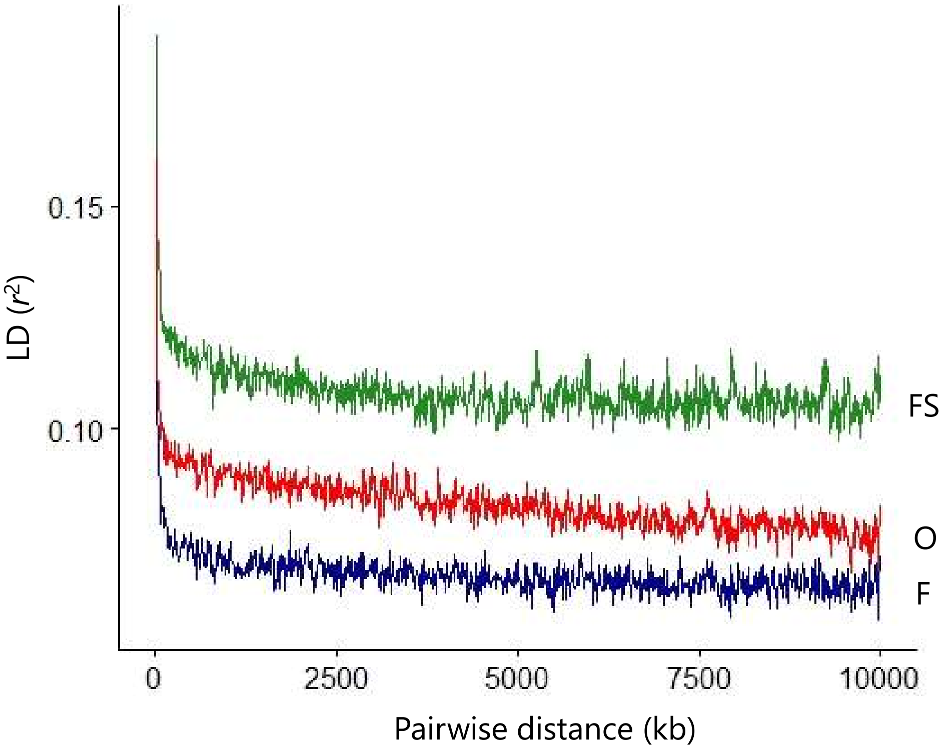


**Fig. S3** Patterns of linkage disequilibrium decay among Japanese cultivars of *Prunus mume*: fruit (F), small-fruit (FS) and ornamental (O) cultivars.

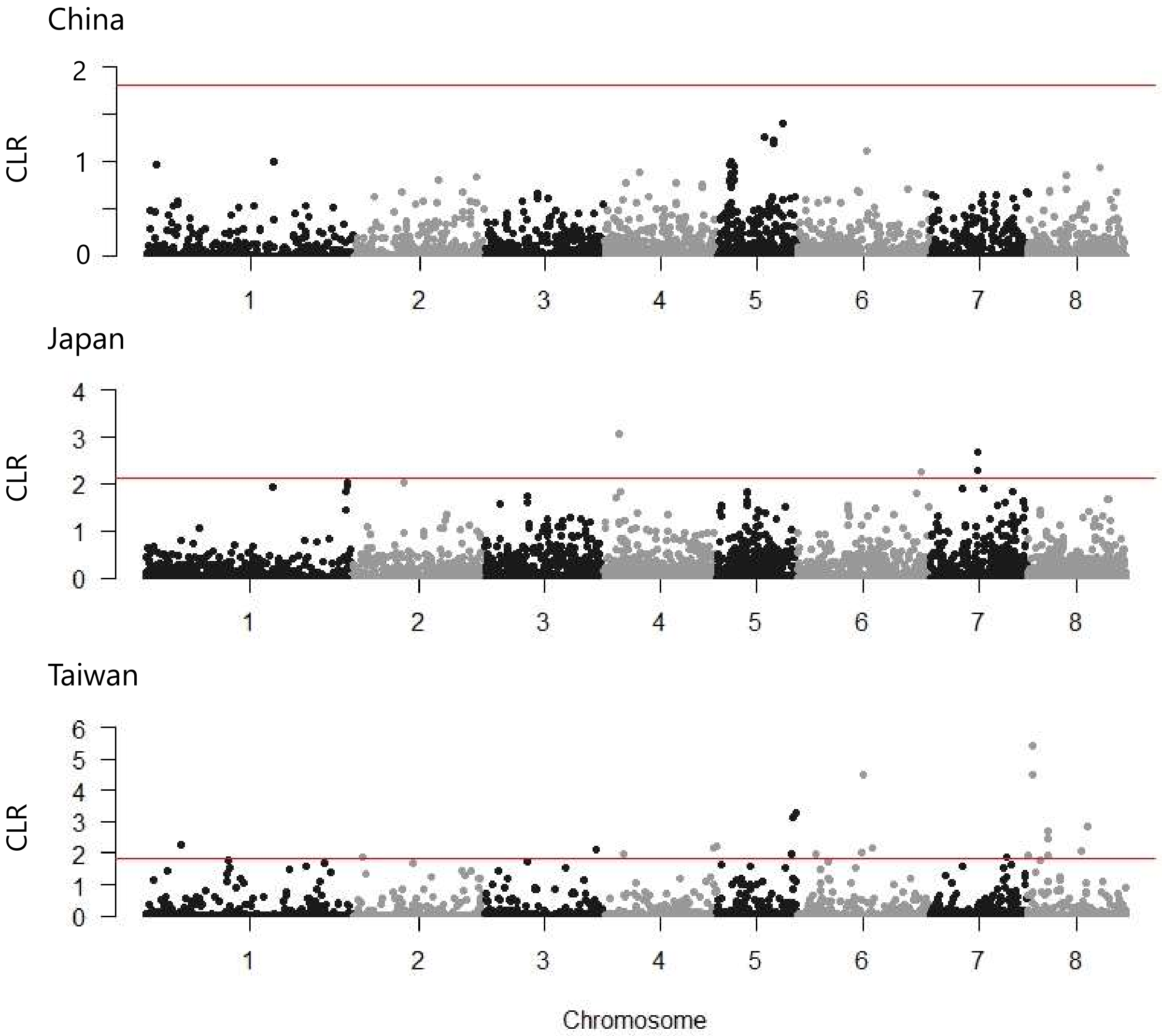


**Fig. S4** Identification of selective sweeps in Chinese, Japanese and Taiwanese cultivars of *Prunus mume* based on site frequency spectrum (SFS)-based SweeD (CLR) analysis. Red bar indicates neutral threshold.

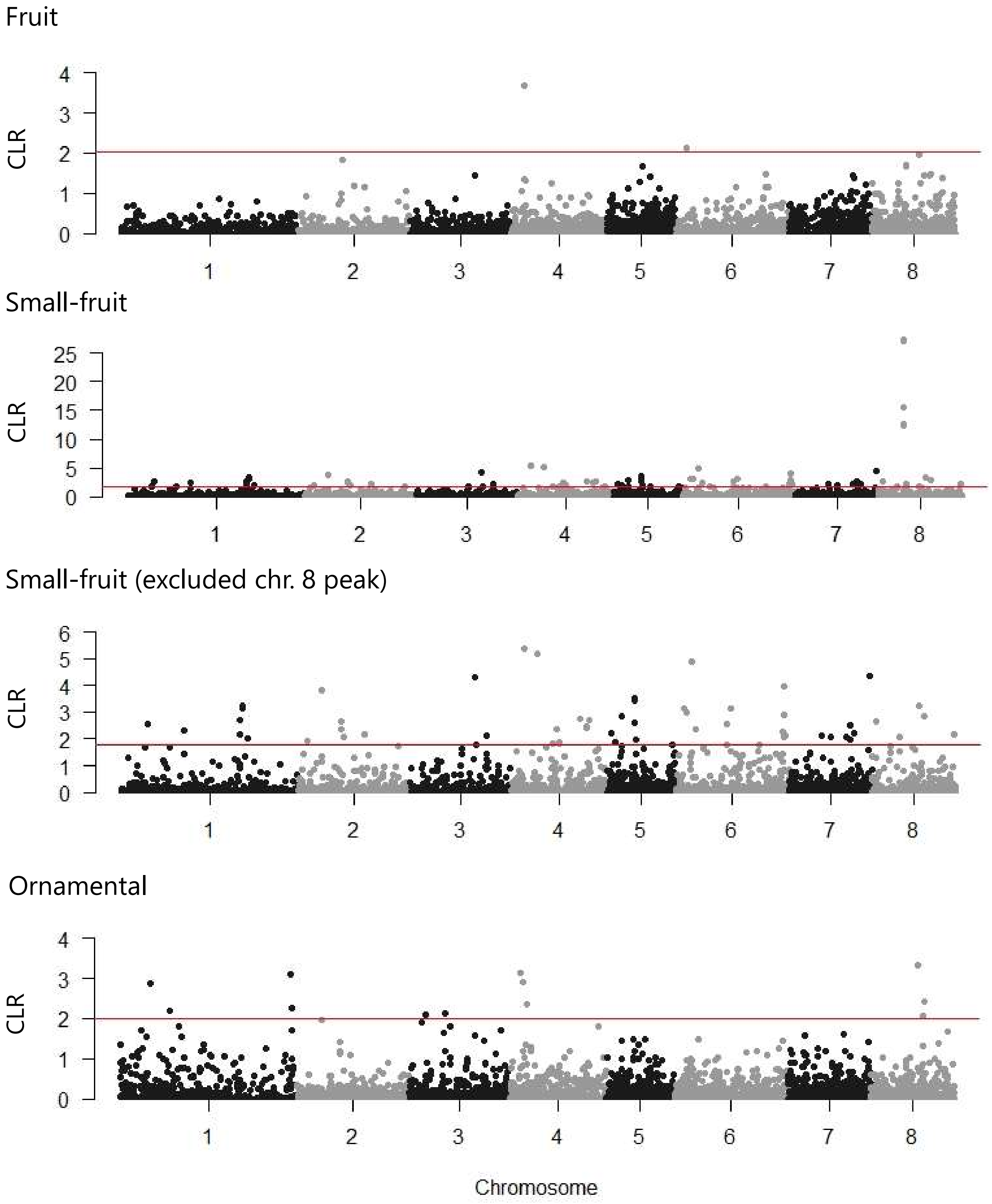


**Fig. S5** Identification of selective sweeps in fruit, small-fruit and ornamental cultivars of *Prunus mume* based on site frequency spectrum (SFS)-based SweeD (CLR) analysis. In small-fruit cultivars, Manhattan plot was redrawn with extremely high peaks in chromosome 8 removed. Red bar indicates neutral threshold.

**(a)** China

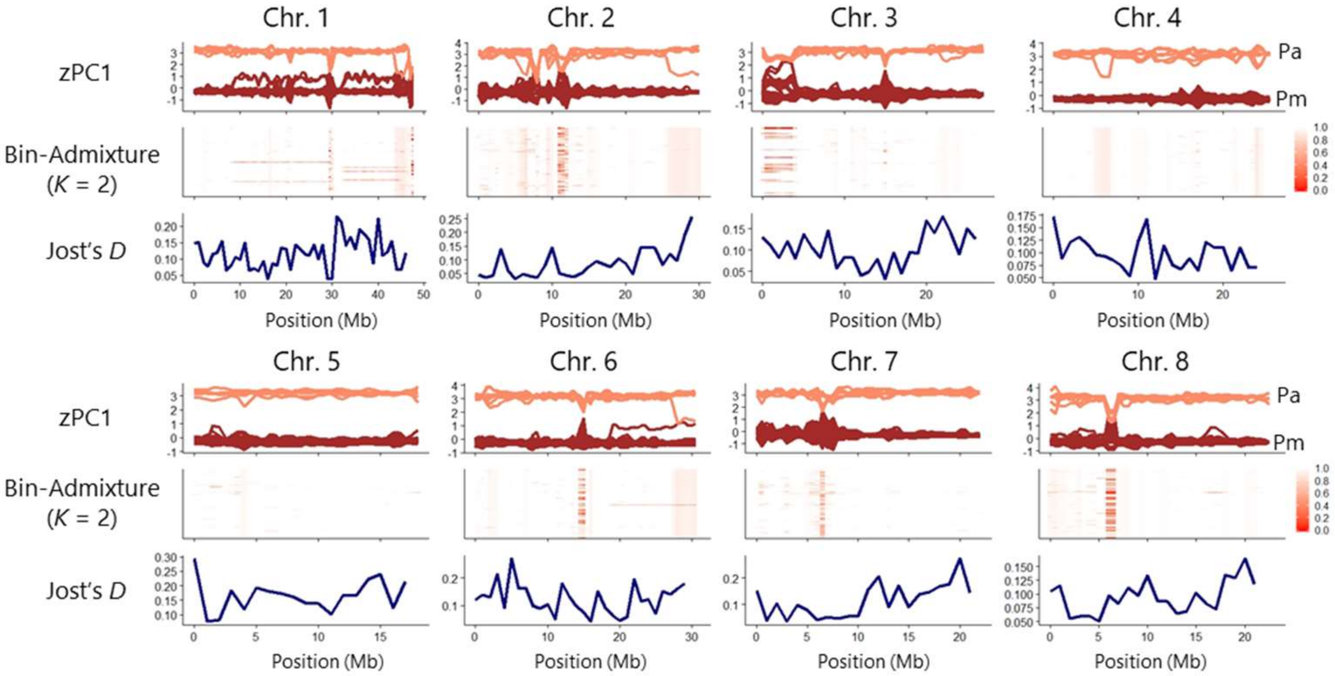


**(b)** Japan

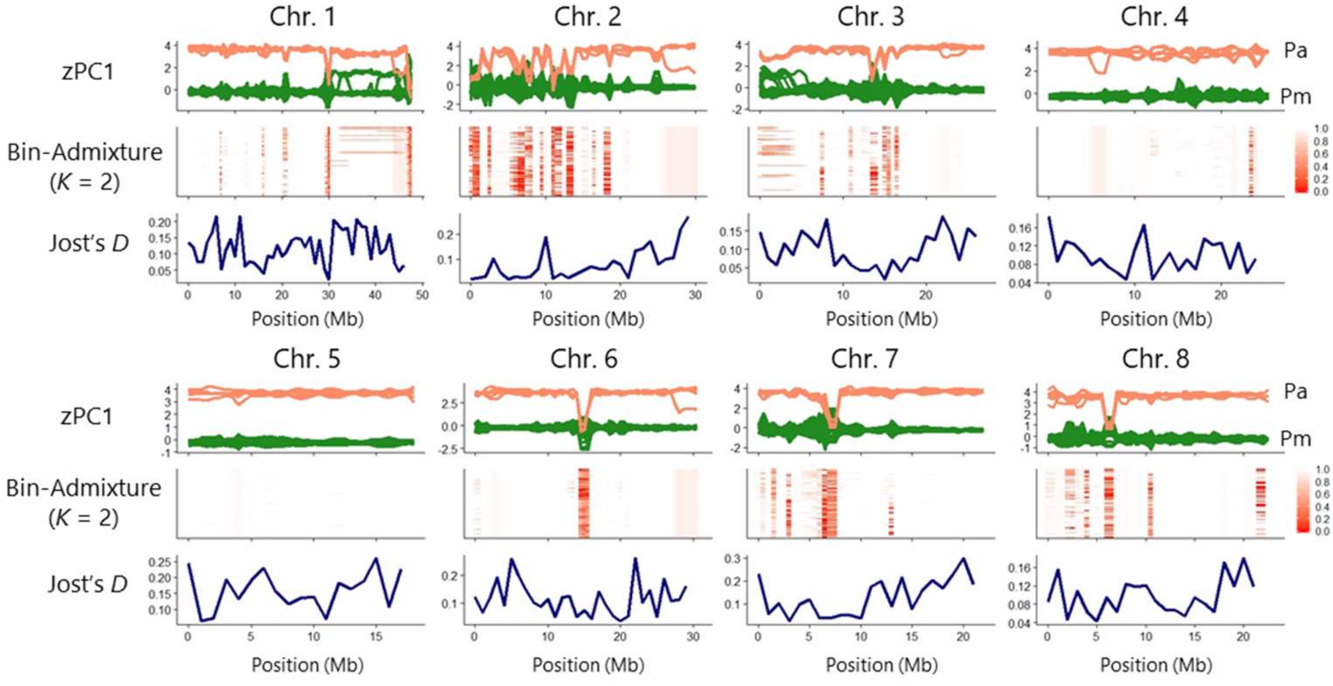


**Fig. S6** Chromosomal patterns of genetic differentiation among, (a) Chinese, (b) Japanese, and (c) Taiwanese cultivars of *Prunus mume* and *P. armeniaca*. All analyses were 1-Mb -binned. zPC1: Z-transformed PC1 calculated with Bin-PCA, Pa: *P. armeniaca*, Pm: *P. mume*. In Bin-Admixture, strength of red color (color scale, 0: highly introgressed–1: no introgression) indicates similarity to *P. armeniaca*.

**(c)** Taiwan

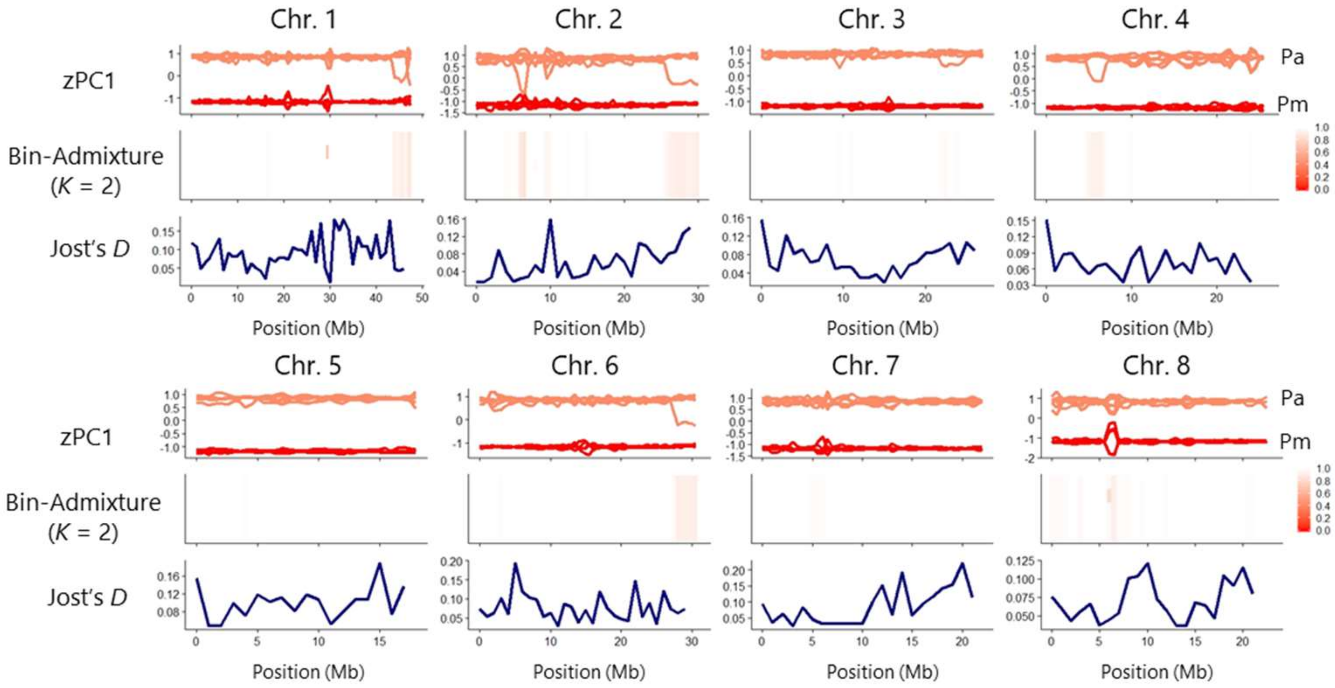


**Fig. S6** Continued.

**(a)** China

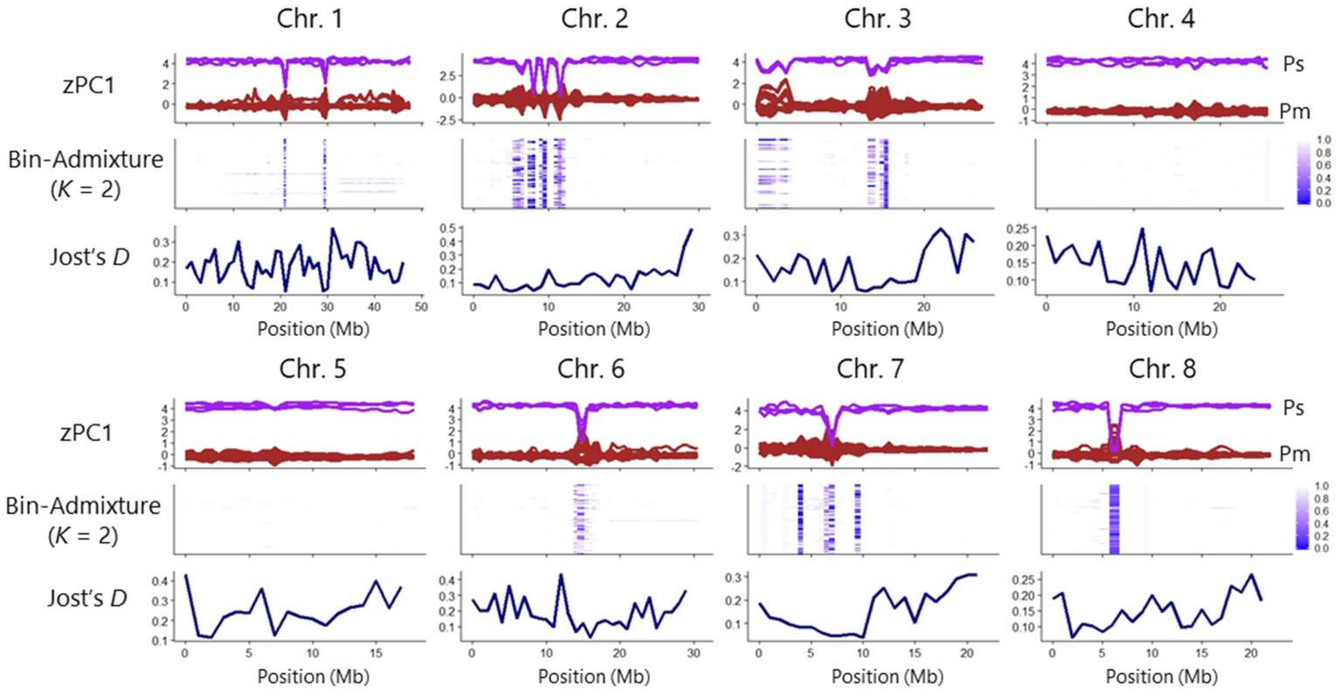


**(b)** Japan

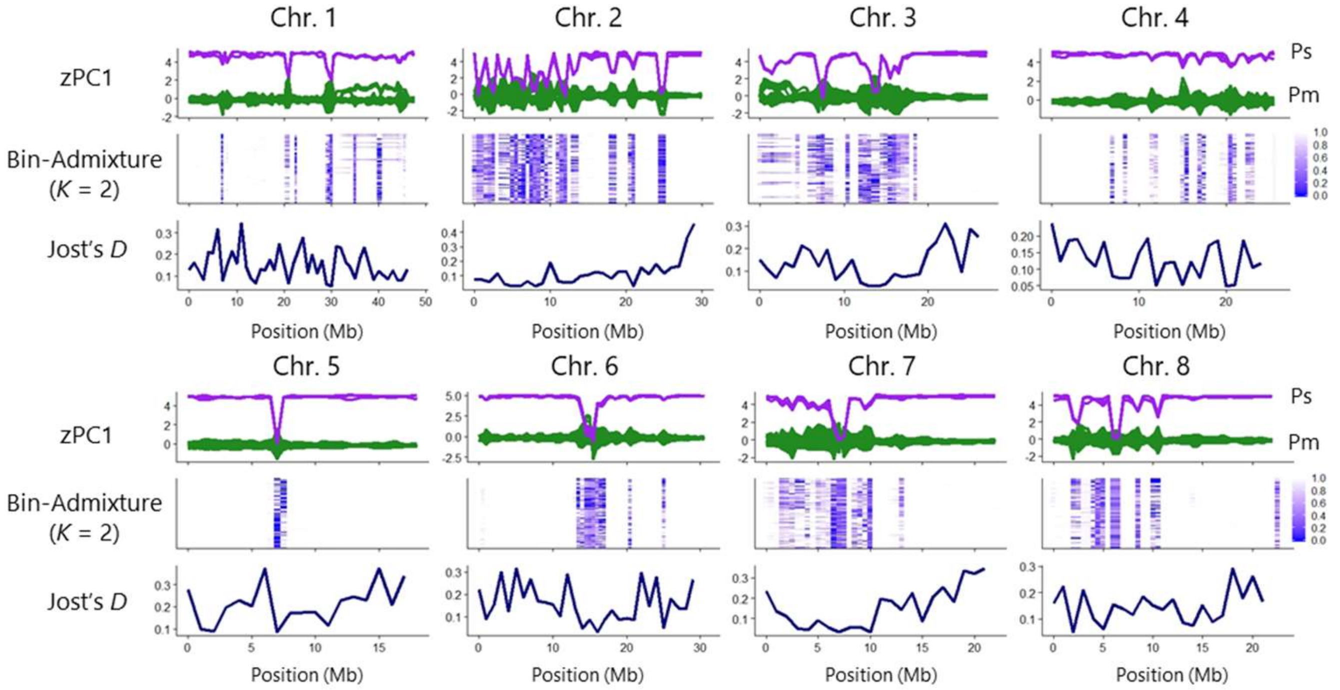


**Fig. S7** Chromosomal patterns of genetic differentiation among, (a) Chinese, (b) Japanese, and (c) Taiwanese cultivars of *Prunus mume* and *P. salicina*. All analyses were 1-Mb -binned. zPC1: Z-transformed PC1 calculated with Bin-PCA, Ps: *P. salicina*, Pm: *P. mume*. In Bin-Admixture, strength of blue color (color scale, 0: highly introgressed–1: no introgression) indicates similarity to *P. salicina*.

**(c)** Taiwan

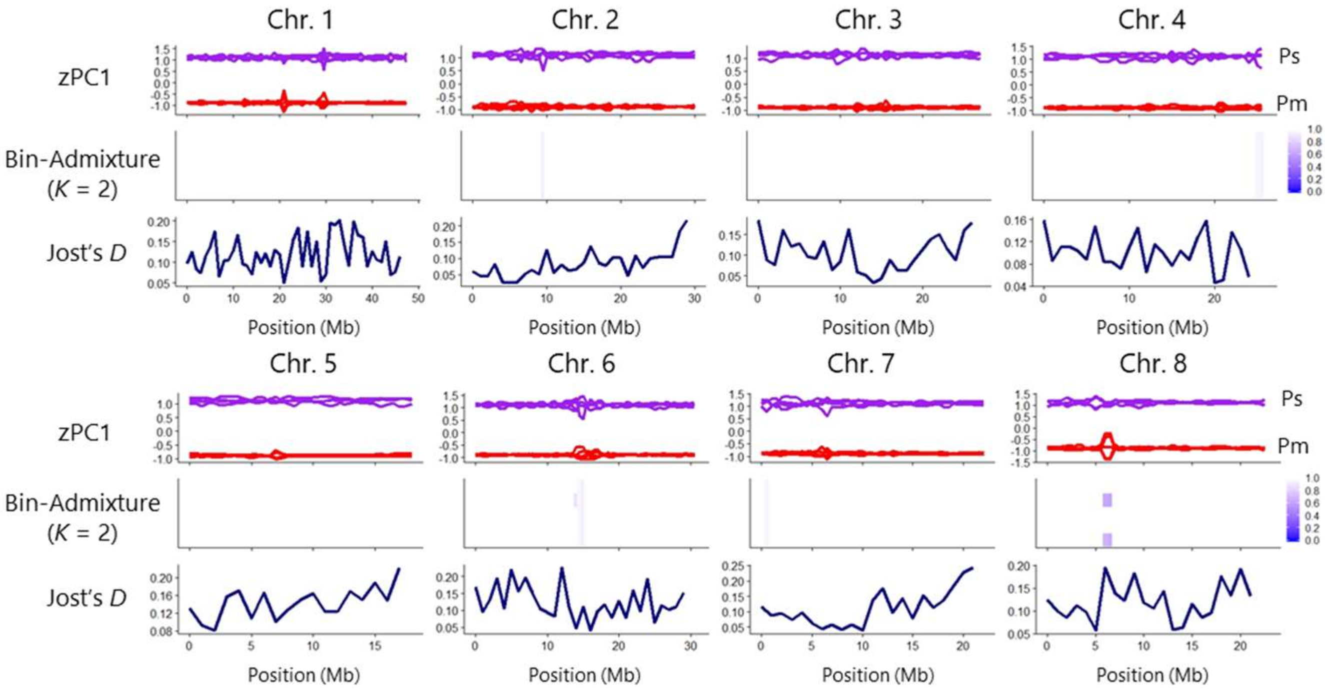


**Fig. S7** Continued.

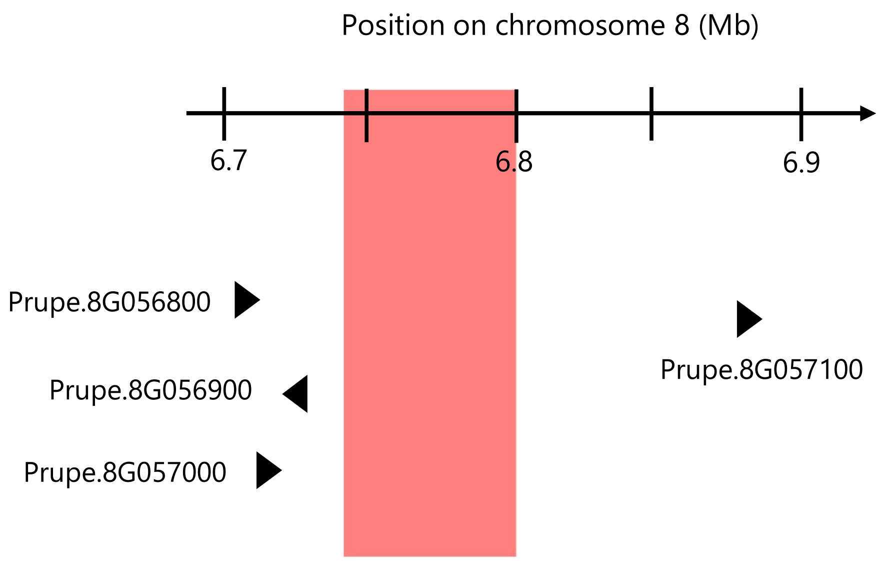


**Fig. S8** Positions of annotated genes adjacent to the candidate region (red) on chromosome 8.

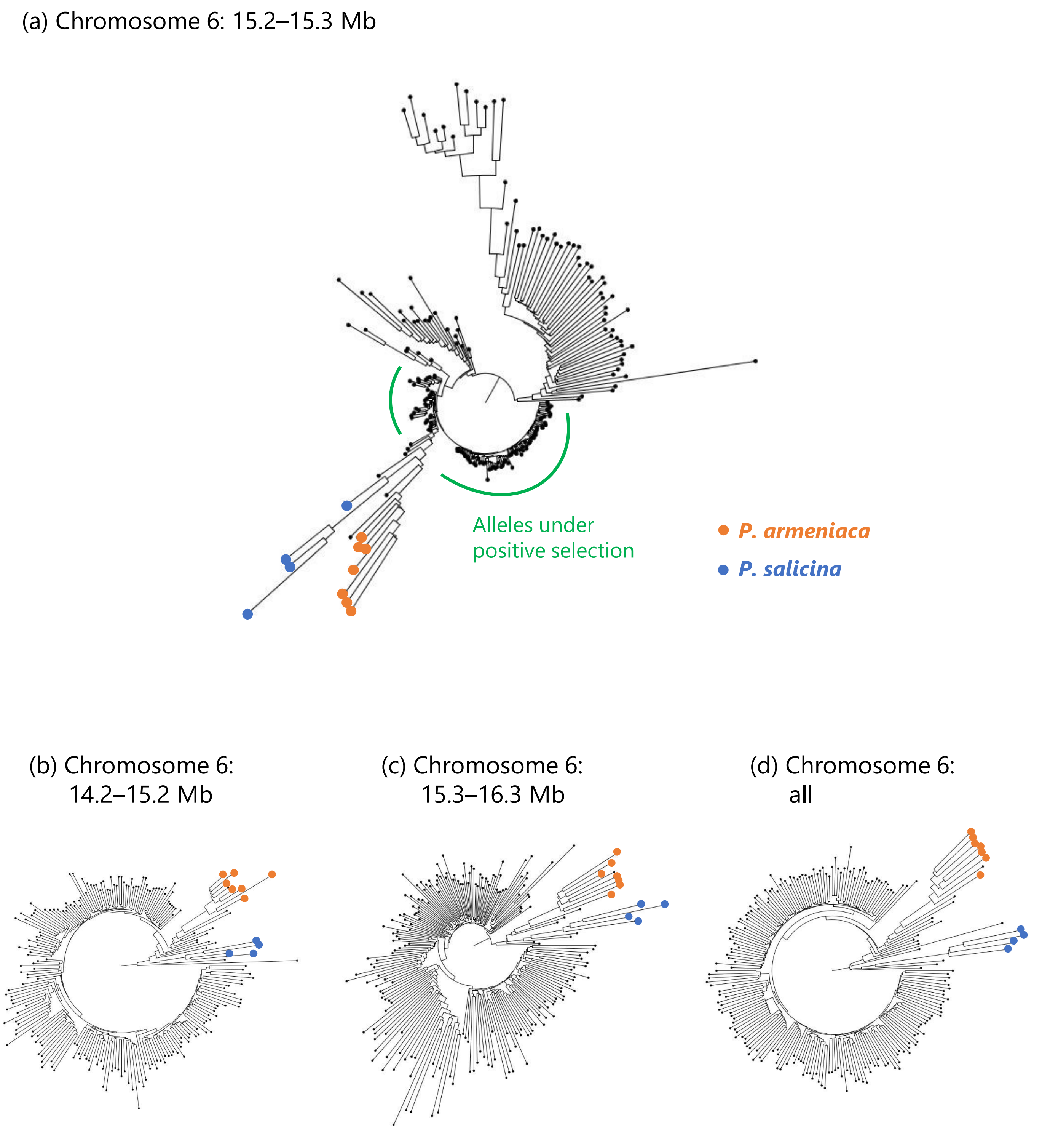


**Fig. S9** Neighbor-joining phylogenetic trees with the single nucleotide polymorphisms (SNPs) in (a) 15.2–15.3, in its (b) upstream and (c) downstream 1-Mb regions and (d) with the whole SNPs in chromosome 6. The tree for the 15.2–15.3-Mb region showed a topology inconsistent with the whole chromosome 6 and the flanking regions (14.2–15.2 and 15.3–16.3 Mb regions), and also inconsistent with the estimated speciation pattern of the subgenus *Prunus*. Alleles which undergone potential selective sweeps were indicated with a green solid line.
